# Sequence-Dependent Conformational Landscapes of Intrinsically Disordered Proteins

**DOI:** 10.1101/2025.08.06.668966

**Authors:** Cong Wang, Bin Zhang

## Abstract

Intrinsically disordered proteins (IDPs) exhibit highly dynamic and heterogeneous conformational ensembles that are strongly influenced by sequence features. While global properties such as chain compaction and scaling behavior have been widely studied, they often fail to resolve the fine-grained, sequence-specific structural variation that underlies IDP function. Here, we perform long-timescale atomistic simulations of 47 representative IDP sequences from the yeast proteome to systematically investigate the relationship between sequence composition and conformational ensemble. To analyze the high-dimensional structural data, we apply uniform manifold approximation and projection (UMAP), a nonlinear dimensionality reduction technique that preserves local structural relationships. The resulting low-dimensional embeddings effectively differentiate IDP ensembles and reveal a novel descriptor—local compactness asymmetry—that quantifies directional differences in chain organization. This metric, denoted *γ_R_*_g_ , captures conformational features orthogonal to traditional global measures such as radius of gyration and end-to-end distance. We show that *γ_R_*_g_ correlates with sequence-level asymmetries in charge and hydropathy, and that conformational dynamics preferentially occur in the more extended region of the chain. The simulation dataset generated in this work also provides a valuable resource for training machine learning models and developing improved coarse-grained force fields for disordered proteins.

## Introduction

Intrinsically disordered proteins (IDPs) are widely present in nature,^1–4^ exhibiting high sequence diversity and playing essential roles in biological processes such as signal transduction, molecular recognition, and regulation.^5–8^ Dysregulation of IDPs can result in a variety of human diseases, including neurodegeneration and cancer.^9–14^ Unlike ordered proteins, IDPs lack a stable, well-defined three-dimensional structure due to their enrichment in hydrophilic amino acids and weak intramolecular interactions.^15–17^ Instead, they adopt dynamic conformations, forming a heterogeneous conformational ensemble that is influenced by both their sequence and environmental conditions. Understanding this ensemble is crucial for elucidating IDPs’ functional mechanisms, characterizing their interactions with other biomolecules, and designing therapeutic strategies targeting disordered regions. ^18,19^

Extensive experimental efforts have been devoted to characterizing the conformational properties of IDPs, using techniques such as nuclear magnetic resonance (NMR) spectroscopy,^20–22^ small-angle X-ray scattering (SAXS),^23,24^ and fluorescence resonance energy transfer (FRET).^25–27^ These methods have provided important insights into the conformational diversity of IDPs. However, their ensemble-averaged nature limits the resolution at which individual conformations can be discerned. Moreover, their inherently low throughput poses practical challenges for large-scale studies aimed at extracting general sequence–ensemble relationships.

Computational modeling offers a valuable complement to experimental approaches by enabling direct access to structural ensembles. Among these, coarse-grained molecular dynamics (MD) simulations are particularly popular due to their efficiency. ^28–34^ Numerous coarse-grained force fields have been developed to simulate IDPs with increasing accuracy, facilitating the rapid generation of conformational ensembles.^35–54^ However, their reduced resolution limits the ability to capture local structural fluctuations and sequence-specific interactions that are likely critical for function. While recent deep learning models have achieved remarkable success in predicting the structure of folded proteins from sequence, ^55,56^ the high conformational heterogeneity of IDPs renders analogous predictions substantially more difficult.^57–60^

Beyond the limitations of current simulation models, there is also a need to revisit the metrics used to describe IDP conformational ensembles. Traditional descriptors—such as radius of gyration, end-to-end distance, asphericity, and the Flory–Huggins scaling exponent—are largely inspired by polymer physics studies of homopolymers.^61–67^ These global measures offer important summary statistics but often fail to capture the sequence-encoded features that distinguish individual IDPs.

In this study, we seek to enrich our understanding of the sequence–ensemble relationship by performing extensive atomistic simulations of IDPs in explicit solvent. Recent advances in force field development have improved the accuracy of atomistic modeling of IDPs, ^68–73^ while modern hardware and software now allow simulations to reach biologically relevant timescales. To systematically explore sequence space, we selected a diverse set of representative IDP sequences from the yeast proteome. The resulting simulation data provide a valuable resource, which we make freely available to the community.

To analyze the high-dimensional conformational ensembles generated by these simulations, we employed uniform manifold approximation and projection (UMAP),^74^ a nonlinear dimensionality reduction technique that preserves local geometric features of the data. By comparing IDP ensembles across different sequences, we identified sequence-dependent structural features not captured by traditional global metrics. In particular, we discovered a novel descriptor that quantifies asymmetric chain compaction, revealing directional preferences in local conformational organization. This analysis also uncovered dynamic correlations between global expansion and local asymmetry, further highlighting the structured nature of IDP fluctuations. Together, these findings provide a refined view of IDP conformational landscapes and demonstrate how high-resolution simulations integrated with data-driven analyses can reveal previously inaccessible features of IDP behavior.

## Results

### Selection of representative intrinsically disordered proteins from the yeast proteome

IDPs exhibit immense diversity in their sequences and compositions, contributing to their varied conformations and functional roles. ^75,76^ A comprehensive characterization of the relationship between sequence and conformational ensemble necessitates a sufficient coverage of the whole sequence space. We select representative sequences from the yeast proteome because of the extensive sequence analysis in prior studies.

Our sequence selection criteria followed the clustering work by Zarin et al. ^77^, with custom modifications to ensure amino acid diversity. Using sequence-based evolutionary features, Zarin et al. ^77^ clustered the IDRs from the *S. cerevisiae* proteome and their orthologous IDRs into 53 groups using a hierarchical clustering algorithm (Figure 1A). These clusters represent IDPs with distinct sequence motifs that encode unique biological functions, providing excellent representation of the diverse biological sequences.

**Figure 1:**
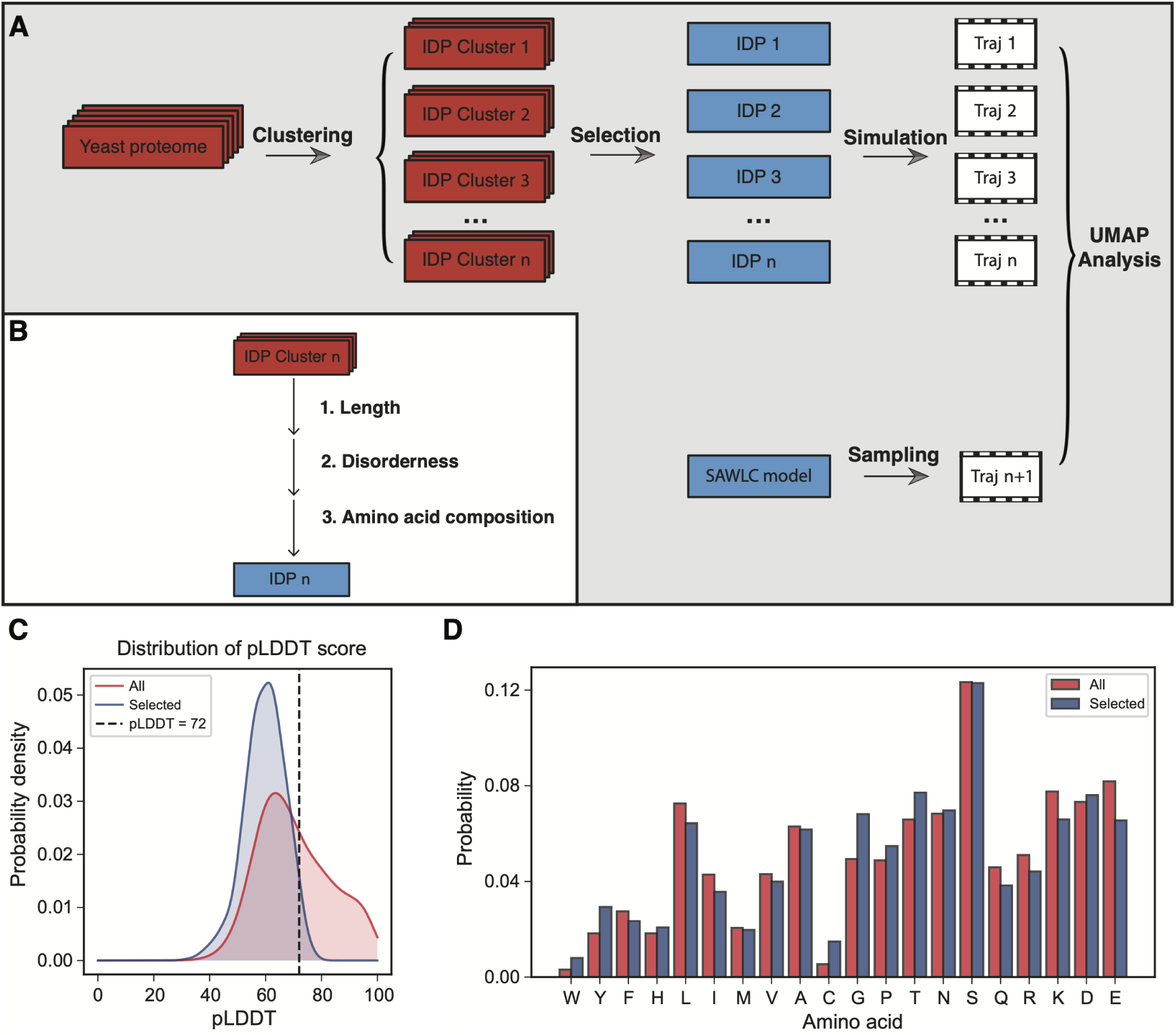
Selection and characterization of representative IDP sequences from the yeast proteome. (A) Schematic overview of the workflow, including sequence clustering, representative selection, atomistic simulations, and ensemble analysis. (B) Criteria used to select one representative 40-residue IDR per cluster, including sequence length, disorder content (pLDDT), and amino acid diversity. (C) Distribution of pLDDT scores across all 40-residue IDPs (red) and the selected representative sequences (blue). The threshold of pLDDT *<* 72, used to define disorder, is indicated. (D) Amino acid composition of all candidate 40-residue IDPs (red) and selected representative sequences (blue), demonstrating that the selected set preserves the overall compositional diversity of the yeast IDR proteome.

We attempted to select one sequence per cluster for further characterization using the following criteria (Figure 1B). First, we restrict the protein sequence length to 40 amino acids, balancing the need to capture essential sequence features and maintaining simulation efficiency. A consistent sequence length further facilitates the comparison of structural ensembles between different IDPs. Secondly, highly disordered sequences were prioritized to minimize the inclusion of ordered regions, which can occasionally be present in IDPs. The degree of disorder was evaluated using per-residue pLDDT scores predicted by ColabFold. ^78^ According to the threshold established by Wilson et al. ^79^, residues with pLDDT scores *<* 72 were classified as disordered. Sequences with more than 90% of disordered residues were retained for further consideration (Figure 1C).

Finally, for the remaining 47 clusters with non-zero sequences, we ranked the sequences in each cluster based on cysteine (C) and tryptophan (W) content. If multiple sequences had identical C and W content, we used amino acid diversity as a secondary criterion, giving priority to sequences with a broader variety of amino acid types. This systematic approach ensured that the top-ranked sequence from each cluster contributed to a balanced and representative amino acid composition across all selected sequences (Figure 1D). The complete list of selected sequences is available in the *Supporting Information*.

### Atomistic simulations generate conformational ensembles for IDPs

We carry out long-time scale simulations of the selected representative sequences using explicit solvent atomistics force fields, a99SB-disp,^80^ which has been shown to accurately capture IDP conformations.^80–83^ Each simulation lasted 10.5 microseconds, a duration sufficient for the IDPs to reach equilibrium (Figure S1).

Consistent with previous studies,^67,86–88^ the Flory–Huggins scaling exponents for most proteins fall within the range of 0.5–0.6 (Figure 2A), indicative of expanded conformations. A small subset of sequences exhibit anomalously low values (below 0.2), which likely reflect numerical artifacts arising from sensitivity in the fitting procedure—particularly due to the short chain length used in this study (see Figure S2). The average radius of gyration (*R*_g_) values also cluster around 1.6 nm (Figure 2D, inset), suggesting a broadly comparable level of compaction across proteins.

**Figure 2:**
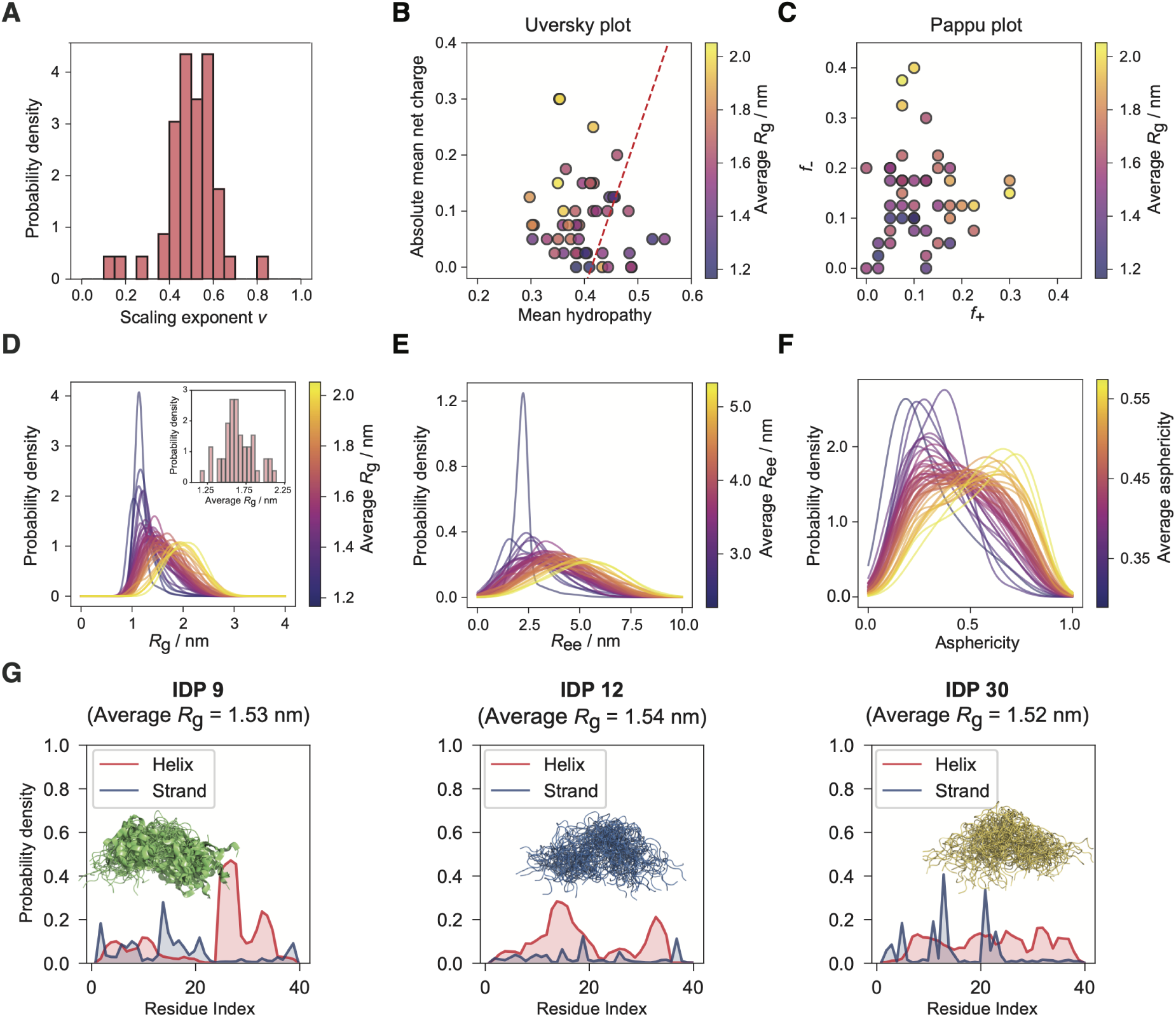
Diverse conformational ensembles revealed by atomistic simulations of distinct IDP sequences. (A) Distribution of Flory–Huggins scaling exponents computed for all simulated IDPs. (B) Scatter plot of mean hydropathy versus absolute mean net charge, with each point representing an IDP sequence and colored by its average *R*_g_. Mean hydropathy is calculated as the average normalized Kyte–Doolittle score^84^ across all residues in the sequence, as introduced by Uversky et al. ^85^. (C) Scatter plot of the fraction of positively charged residues versus the fraction of negatively charged residues for each IDP sequence, with points colored by average *R*_g_ (Pappu plot). (D) Probability density distributions of *R*_g_ for each IDP ensemble, with curves colored by their respective average *R*_g_. The inset shows the distribution of average *R*_g_ values across all sequences. (E) Probability density distributions of end-to-end distance (*R*_ee_) for each IDP ensemble, colored by their average *R*_ee_. (F) Probability density distributions of asphericity for each ensemble, with curves colored by the corresponding average asphericity. (G) Representative IDPs with nearly identical average *R*_g_ values, highlighting their distinct local secondary structure propensities along the sequence. Line plots show residue-wise probabilities of helix and strand formation.

The distributions of *R*_g_, end-to-end distance (*R*_ee_), and asphericity further reinforce this observation, showing substantial overlap across different IDP ensembles (Figures 2D–F and Figures S3–S5). These similarities imply a degree of global conformational resemblance when assessed using conventional shape descriptors.

This resemblance is in line with sequence-level features known to influence protein size. As shown in Figure 2B, most IDPs share moderate mean hydropathy and net charge, conditions associated with extended ensembles. IDPs exhibiting larger average *R*_g_ values tend to deviate from this trend, typically displaying lower hydropathy and higher net charge. A similar pattern emerges in the Pappu plot (Figure 2C), where most IDPs show comparable fractions of positively and negatively charged residues, while sequences with more asymmetric charge distributions tend to have larger *R*_g_ values.

However, visual inspection of representative 3D structures reveals substantial differences (Figure S6), even among IDPs with nearly identical *R*_g_ values (Figure 2G). This diversity is further reflected in the residue-wise probability of secondary structure formation (Figure 2G and S7), and in the distribution of root-mean-square deviations (RMSD) relative to IDP 1 (Figure S8).

Together, these results underscore the limitations of global descriptors such as *R*_g_ in capturing critical, sequence-specific conformational features of IDPs. While secondary structure profiles can reveal distinctions between ensembles, their discrete nature makes them less convenient for detailed structural analyses. This motivates the development of continuous, numeric metrics that can more effectively characterize and differentiate IDP conformations.

### UMAP embeddings differentiate IDP conformational landscapes

To systematically distinguish IDP conformations and identify new structural descriptors, we applied Uniform Manifold Approximation and Projection (UMAP) analysis.^74^ As described in the Methods section, UMAP is a dimensionality reduction technique that generates low-dimensional projections that best preserve the topological structure of the original highdimensional data.^74^ This approach maintains local neighborhood relationships—conformations that are similar in the high-dimensional space remain close in the projection—while also ensuring that structurally divergent conformations are well separated. This allows the resulting low-dimensional embeddings to effectively consolidate structural information and accentuate key differences across the ensemble of 3D conformations.

Each conformation, sampled from atomistic simulations, was represented as a vector encoding the non-redundant pairwise distances between all *α*-carbon atoms. To provide a reference for comparison, we also included conformations generated from the self avoiding worm like chain (SAWLC) polymer model^89^ (see Methods). These vectors served as input to the UMAP algorithm, which mapped the high-dimensional structural data onto two variables, denoted as UMAP1 and UMAP2, collectively referred to as the UMAP embedding.

As shown in Figure S9, the UMAP embeddings of different IDP sequences form distinct distributions, underscoring the sequence dependence of IDP conformational ensembles. To illustrate the utility of these embeddings, we focus on IDPs 9, 12, and 30, which differ markedly in sequence and secondary structure content (Figure 3A) but exhibit nearly identical average *R*_g_ values and similar *R*_g_ distributions (Figure 3B). Despite these similarities in global metrics, their UMAP embeddings are clearly separated (Figure 3C), particularly along the UMAP2 axis (Figure 3D), indicating differences in their dominant conformational states. This demonstrates that UMAP effectively resolves sequence-specific conformational features that are not captured by *R*_g_ alone.

**Figure 3:**
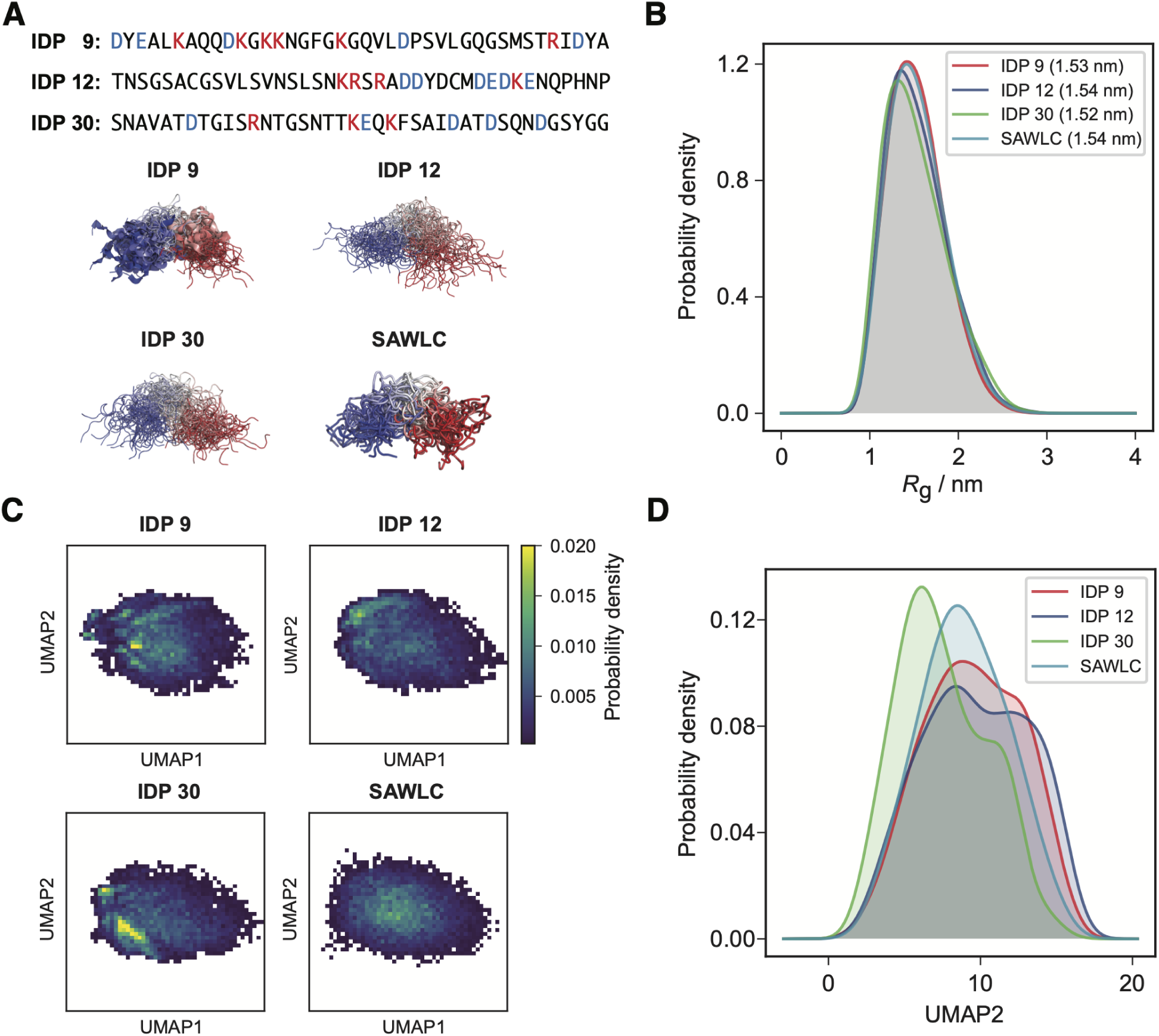
UMAP embeddings differentiate IDP conformational landscapes. (A) Top: Amino acid sequences of IDPs 9, 12, and 30, with positively charged residues shown in red and negatively charged residues in blue. Bottom: Representative conformations from atomistic simulations of the three IDPs, and for the SAWLC model. Structures are colored from blue (N-terminus) to red (C-terminus). (B) Probability density distributions of *R*_g_ for the four ensembles, illustrating their similar global compaction. (C) Two-dimensional probability density distributions of UMAP embeddings for the same ensembles, showing clear separation in conformational space despite similar *R*_g_. (D) One-dimensional probability density distributions along the UMAP2 axis, highlighting differences in dominant conformational features not resolved by global metrics.

Although the UMAP1 and UMAP2 distributions peak at different values for each protein, the ranges of the embeddings exhibit considerable overlap. This reflects the flexible nature of IDPs, whose conformational states form a continuum rather than discrete structural classes. This pattern holds across all simulated IDPs: the overall distribution of UMAP embeddings spans a continuous space with no sharply defined clusters (Figure S10), consistent with the inherent heterogeneity and dynamic nature of IDP ensembles.

To evaluate the robustness of our approach, we also tested an alternative representation for UMAP input: the RMSD of each conformation relative to a set of reference structures. The resulting embeddings closely resemble those obtained from pairwise *α*-carbon distances (Figure S11A), further supporting the reliability and consistency of the analysis.

### Linking UMAP Embeddings to Structural and Sequence Features

While the UMAP embeddings effectively differentiate IDP conformations, their physical interpretation is not immediately apparent. To better understand the structural features encoded by each axis, we examined representative conformations along UMAP1 and UMAP2. As shown in Figure 4A, structures evolve smoothly from compact to extended as UMAP1 increases, indicating that this axis primarily reflects changes in global size. In contrast, UMAP2 captures asymmetry in chain compaction—specifically, how structural compactness is distributed between the two termini—revealing features not accessible through conventional global metrics.

**Figure 4:**
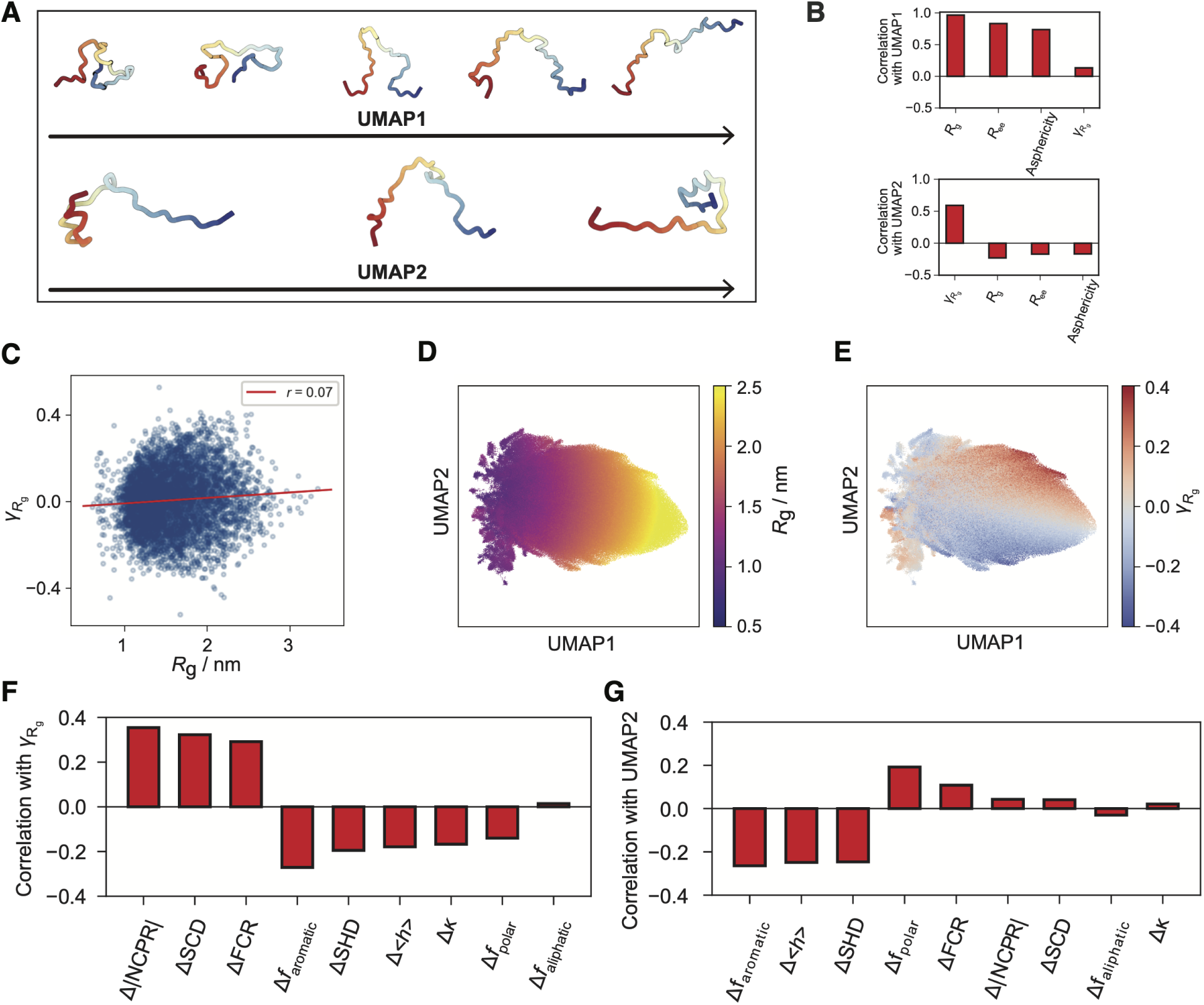
Linking UMAP embeddings to structural and sequence Features. (A) Representative conformations illustrating structural variation along the UMAP1 and UMAP2 axes. (B) Pearson correlation coefficients between UMAP1 or UMAP2 and various conformational descriptors, including *R*_g_, *R*_ee_, asphericity, and local compactness asymmetry (*γ_R_*_g_ ). (C) Scatter plot of radius of gyration (*R*_g_) versus *γ_R_*_g_ , illustrating their low correlation. (D) UMAP embedding of all conformational ensembles, with each point colored by *R*_g_. (E) UMAP embedding of all conformational ensembles, with each point colored by *γ_R_*_g_. (F) Correlation between *γ_R_*_g_ and differences in sequence features between the N- and C-terminal halves of each IDP. (G) Correlation between UMAP2 values and the same sequence feature differences as in (F).

To provide a more quantitative interpretation of the UMAP embeddings, we evaluated their correlations with several commonly used structural descriptors: the radius of gyration (*R*_g_), end-to-end distance (*R*_ee_), asphericity, and a newly defined metric, *γ_R_*_g_ . The latter quantifies local compactness asymmetry along the chain and is defined as

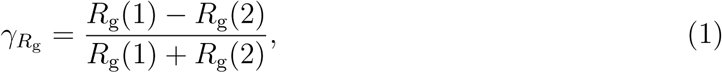

where *R*_g_(1) and *R*_g_(2) are the radii of gyration computed from the first and second halves of the sequence, respectively.

As summarized in Figure 4B and Figures S12–S13, UMAP1 exhibits strong correlations with *R*_g_, *R*_ee_, and asphericity, with the highest correlation observed for *R*_g_. This confirms that UMAP1 retains key global size and shape features after dimensionality reduction. In contrast, UMAP2 shows weak correlations with these conventional descriptors but is strongly correlated with *γ_R_*_g_ , suggesting that UMAP2 captures orthogonal structural variation related to local asymmetry. These relationships are further illustrated in Figures 4D, 4E, S14 and S15.

Notably, *γ_R_*_g_ is nearly uncorrelated with *R*_g_ itself (Figure 4C), whereas *R*_ee_ and asphericity exhibit strong correlations with *R*_g_ (Figure S16), highlighting *γ_R_*_g_ as a complementary descriptor. While global metrics characterize overall compactness, *γ_R_*_g_ resolves asymmetry along the sequence, providing a more nuanced view of IDP structural variability.

To investigate the sequence determinants of this asymmetry, we computed the correlation between *γ_R_*_g_ and the difference in several common sequence features across the N- and C-terminal halves of each chain (Figure 4F). For a given sequence feature *f* , the difference is defined as

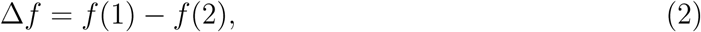

where *f* (1) and *f* (2) are the values of *f* computed over the first and second halves of the sequence, respectively.

The sequence features analyzed include the absolute net charge per residue (|NCPR|), sequence charge decoration (SCD),^65^ fraction of charged residues (FCR), fraction of aromatic residues (*f*_aromatic_), sequence hydropathy difference (SHD),^65^ mean hydropathy (⟨*h*⟩), *κ* parameter,^62^ and the fractions of polar and aliphatic residues (*f*_polar_ and *f*_aliphatic_, respectively). Among these, Δ|NCPR| exhibits the strongest correlation with *γ_R_*_g_ , indicating that asymmetry in charge density along the chain is a key determinant of local compactness differences. Specifically, a higher |NCPR| in the N-terminal half or a lower |NCPR| in the C-terminal half is associated with a larger *γ_R_*_g_ , corresponding to greater extension of the N-terminal segment.

Other sequence features also show non-zero correlations with *γ_R_*_g_ , though none exceed an absolute value of 0.4, suggesting that *γ_R_*_g_ reflects a composite effect of multiple sequence characteristics rather than being dominated by a single factor. These results highlight the multifactorial nature of sequence-dependent conformational asymmetry in IDPs. We observe similar trends in the correlation between UMAP2 and these same sequence feature differences (Figure 4G), further supporting the interpretation that UMAP2 captures local structural asymmetry and its underlying sequence determinants.

### Sequence-encoded bias in IDP conformational fluctuations

The pronounced asymmetry observed at the termini of various IDP conformations prompted us to further investigate their underlying dynamics. We hypothesize that the conventional coil–globule transition model, ^90^ which resembles a first-order phase transition, may be insufficient to describe the conformational behavior of IDPs. Classical coil–globule transitions typically assume an isotropic, global collapse of the chain without directional bias. In contrast, the variation observed along UMAP2 and the *γ_R_*_g_ metric suggests a potential preference for asymmetric compaction—where one end of the chain may collapse more readily than the other. This directional bias implies that IDP dynamics may deviate from the uniform behavior predicted by standard polymer models and instead reflect sequence-encoded preferences for localized structural reorganization.

To investigate the coupling between global and local conformational dynamics in IDPs, we computed the time-lagged cross-correlation between fluctuations in radius of gyration (*R*_g_) and local compactness asymmetry (*γ_R_*_g_ ):

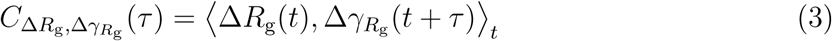

where Δ*R*_g_(*t*) = *R*_g_(*t* + 1, ns) − *R*_g_(*t*), and Δ*γ_R_*_g_ (*t*) is defined analogously. Figure 5A (and Figure S17) summarizes *C*(*τ* ) for all IDP systems alongside the SAWLC polymer model as a reference.

**Figure 5:**
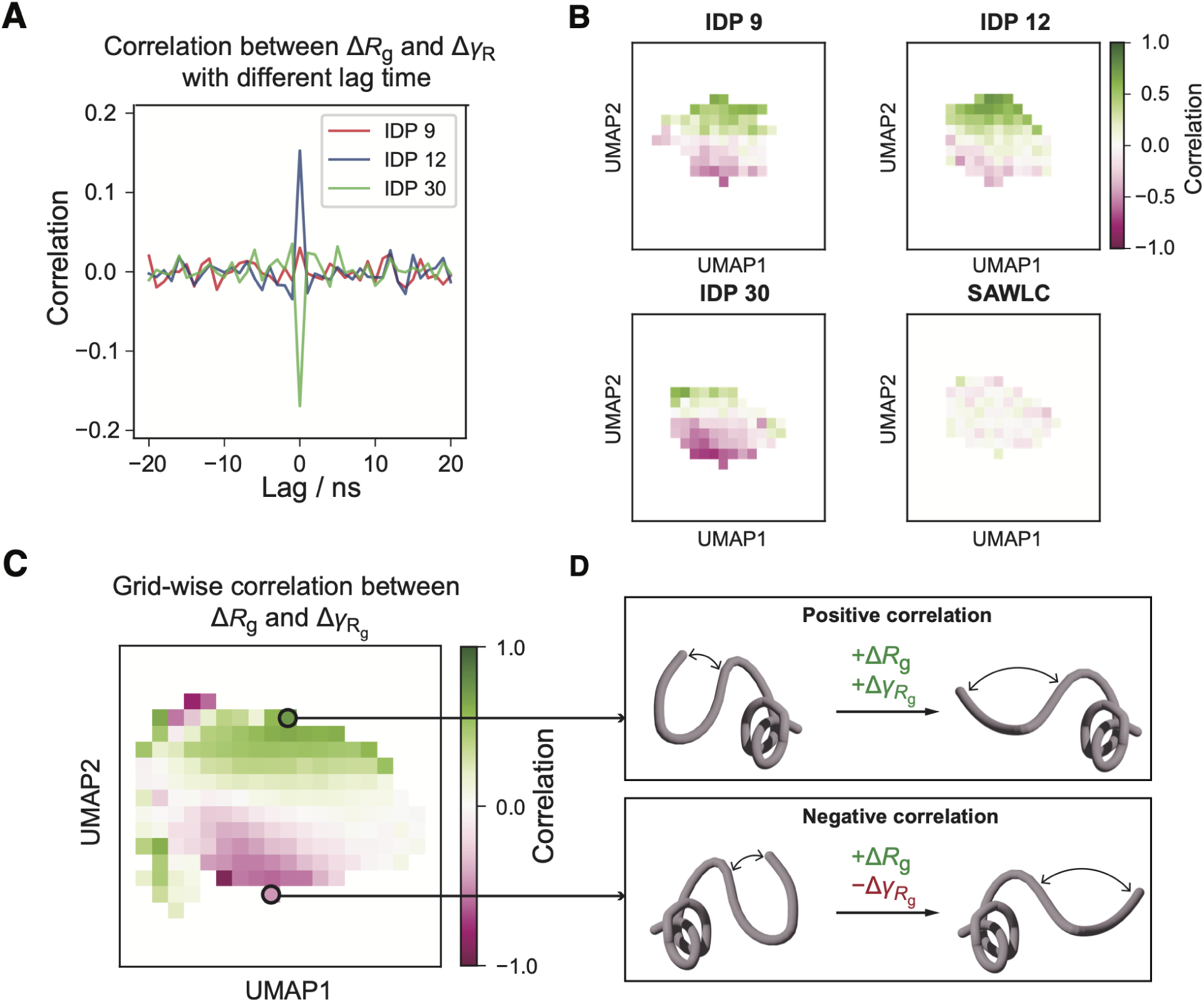
Dynamic coupling between global expansion and local asymmetry in IDP conformations. (A) Time-lagged cross-correlation between fluctuations in Δ*R*_g_(*t*) and local compactness asymmetry Δ*γ_R_*_g_ (*t*) for three representative IDP ensembles. (B) Grid-wise cross-correlation at zero lag time (*τ* = 0) between Δ*R*_g_(*t*) and Δ*γ_R_*_g_ (*t*) mapped onto the UMAP embeddings of IDPs 9, 12, and 30, and the SAWLC model. (C) Zero-lag cross-correlation between Δ*R*_g_(*t*) and Δ*γ_R_*_g_ (*t*) across all IDP ensembles, visualized over the UMAP embedding. (D) Schematic illustrations of positive and negative coupling between Δ*R*_g_ and Δ*γ_R_*_g_ , corresponding to extended N-terminal or C-terminal regions, respectively, highlighting directional bias captured by UMAP2.

Despite exhibiting similar *R*_g_ distributions, the IDPs (e.g., IDP 9, 12, and 30) display distinct correlation profiles, indicating heterogeneity in their conformational dynamics. In particular, the zero-lag correlation *C*(0) varies substantially across systems—near zero for IDP 9, but positive or negative for IDPs 12 and 30, respectively. These correlations decay rapidly with increasing lag, typically vanishing beyond 1 ns, consistent with the fast internal dynamics of disordered proteins.

To further resolve the spatial origin of this dynamic coupling, we mapped *C*(0) values across the UMAP embedding (Figure 5B and Figure S18). As expected, the SAWLC ensemble displays negligible correlations throughout the landscape, consistent with its lack of structured dynamics. In contrast, the IDP ensembles exhibit a clear spatial trend: conformations with large UMAP2 values (corresponding to *γ_R_*_g_ *>* 0) are enriched in positive *C*(0) values, whereas those with negative *γ_R_*_g_ show negative correlations. These observations indicate that the sign and magnitude of dynamic coupling between global and local shape fluctuations are linked to specific regions of conformational space.

This trend can be rationalized by considering the physical interpretation of *γ_R_*_g_ . When *γ_R_*_g_ *>* 0, the N-terminal half of the chain is more extended than the C-terminal half. In this case, increases in *R*_g_ are primarily driven by fluctuations in the already extended N-terminal region, while the compact C-terminal region remains relatively unchanged—resulting in a positive correlation between Δ*R*_g_ and Δ*γ_R_*_g_ . The opposite scenario occurs when *γ_R_*_g_ *<* 0: fluctuations in *R*_g_ arise from the extended C-terminal segment, leading to negative correlations. Despite the sign reversal, both cases point to a shared physical mechanism: dynamic fluctuations preferentially originate from the more extended, and therefore more flexible, region of the chain. Compact regions, stabilized by local interactions, contribute less to the overall conformational change.

## Conclusions and Discussion

In this study, we employed atomistic simulations to generate conformational ensembles for representative IDP sequences selected from the yeast proteome. To analyze the resulting high-dimensional structural data, we applied UMAP for dimensionality reduction, which revealed the heterogeneous nature of IDP ensembles and enabled systematic comparisons across sequences. Through examining the physical interpretation of the UMAP embeddings, we identified a new descriptor, local compactness asymmetry (*γ_R_*_g_ ), which captures sequence-dependent structural features not reflected in conventional global conformational metrics such as radius of gyration and end-to-end distance. This descriptor provides complementary insight by quantifying asymmetric chain compaction within IDP ensembles, thereby uncovering structural variations that are not apparent from global shape alone. Moreover, we demonstrated that *γ_R_*_g_ correlates with sequence features, including properties and residue hydrophobicities, highlighting the influence of sequence features on local conformational organization. These results establish a framework for integrating atomistic simulations with dimensionality reduction to investigate sequence-encoded structural diversity in disordered proteins.

Beyond its immediate use in revealing sequence-dependent conformational features, the dataset generated in this study provides a valuable resource for the broader community. The large collection of atomistic simulations across diverse IDP sequences offers a high-resolution benchmark for investigating sequence-to-ensemble relationships. These data can be directly leveraged to train or evaluate machine learning models aimed at predicting structural ensembles from sequence, particularly those employing deep generative frameworks.^53,57–60,91^ Furthermore, the conformational ensembles serve as a rich training set for parameterizing and validating coarse-grained force fields, enabling the development of transferable models that more accurately capture IDP behavior across different sequence and environmental contexts.^33^

## Methods

### Explicit solvent atomistic simulations

We performed explicit-solvent atomistic simulations for each individual IDP as follows. Initial structures were generated using ColabFold, ^78^ and the system setup, including coordinate, topology, and force field files, was carried out using GROMACS.^92^ Each IDP was solvated in a dodecahedral water box with a box length of 8.0 nm, and NaCl was added to a final concentration of 0.15 M. Simulations employed the a99SB-*disp* force field for proteins, along with its corresponding water model.^80^

Molecular dynamics simulations were executed using the OpenMM package. ^93^ After energy minimization, production simulations were run in the constant pressure, and temperature (NPT) ensemble at 300 K and 1 bar for 10.5 *µ*s. We applied the hydrogen mass repartitioning scheme^94^ to enable a 4 fs integration timestep and used the Langevin middle integrator^95^ for temperature control. Configurations were saved every 100 ps, yielding 105,000 frames per trajectory for each of the 47 simulated IDPs.

To verify the adequacy of the simulation box, we monitored the minimum image distances between atoms across periodic boundaries; no artifacts from inter-box interactions were observed (Figure S19).

### Self avoiding worm like chain (SAWLC) model

The SAWLC model was used to generate a reference ensemble for comparison with ensembles obtained from atomistic simulations. In this model, the number of beads and bond length were set to 40 and 0.38 nm, respectively. The model incorporated a persistence length of 0.4 nm and a linker diameter of 0.38 nm. SAWLC ensembles were generated using the PolymerCpp Python package,^96^ with a total of 105,000 samples produced.

### Conformational ensemble analysis

We applied several metrics to characterize IDP conformations: the radius of gyration (*R*_g_), the end-to-end distance (*R*_ee_), the asphericity, the radius-of-gyration symmetry ratio (*γ_R_*_g_), and the Flory–Huggins scaling exponent. The values of *R*_g_, *R*_ee_, and *γ_R_*_g_ were obtained using the Python package MDTraj, ^97^ while the asphericity and the Flory–Huggins scaling exponent and corresponding reduced *χ*^2^ test value were calculated using the Python package SOURSOP.^98^

### UMAP analysis

We applied Uniform Manifold Approximation and Projection (UMAP)^74^ to perform dimensionality reduction across all 47 IDP ensembles, along with a reference ensemble generated from the SAWLC polymer model, enabling direct comparisons across systems. To reduce computational cost, we subsampled each trajectory by retaining every 10th frame, resulting in 10,500 conformations per ensemble.

Two types of structural descriptors were used as input to the UMAP algorithm: (1) non-redundant pairwise distances between all *α*-carbon atoms in each conformation, and (2) root-mean-square deviation (RMSD) of each conformation relative to a common set of reference structures. Reference conformations were uniformly selected every 2,000 frames across all ensembles to ensure a consistent comparison basis.

Unless stated otherwise, the pairwise distance descriptor was used for all figures in the main text. UMAP parameters were set with n neighbors = 200 and min dist = 0.99. All other settings were kept at their default values.

## Acknowledgement

This work was supported by the National Institutes of Health (Grant R35GM133580).

## Data and materials availability

MD simulation trajectories of IDPs and the structural ensemble generated using the SAWLC model are available at [Zenodo](https://zenodo.org/records/15493750).

## References

(1) Dunker, A. K.; Romero, P.; Obradovic, Z.; Garner, E. C.; Brown, C. J. Intrinsic protein disorder in complete genomes. Genome informatics 2000, 11, 161–171.

(2) Ward, J. J.; Sodhi, J. S.; McGuffin, L. J.; Buxton, B. F.; Jones, D. T. Prediction and functional analysis of native disorder in proteins from the three kingdoms of life. Journal of molecular biology 2004, 337, 635–645.

(3) Pancsa, R.; Tompa, P. Structural disorder in eukaryotes. PloS one 2012, 7, e34687.

(4) Latham, A. P.; Zhang, B. Molecular Determinants for the Layering and Coarsening of Biological Condensates. Aggregate 2022, 3, 1–9.

(5) Xie, H.; Vucetic, S.; Iakoucheva, L. M.; Oldfield, C. J.; Dunker, A. K.; Uversky, V. N.; Obradovic, Z. Functional anthology of intrinsic disorder. 1. Biological processes and functions of proteins with long disordered regions. Journal of proteome research 2007, 6, 1882–1898.

(6) Wright, P. E.; Dyson, H. J. Intrinsically disordered proteins in cellular signalling and regulation. Nature reviews Molecular cell biology 2015, 16, 18–29.

(7) Bah, A.; Forman-Kay, J. D. Modulation of intrinsically disordered protein function by post-translational modifications. Journal of Biological Chemistry 2016, 291, 6696–6705.

(8) Tantos, A.; Han, K.-H.; Tompa, P. Intrinsic disorder in cell signaling and gene transcription. Molecular and cellular endocrinology 2012, 348, 457–465.

(9) Vavouri, T.; Semple, J. I.; Garcia-Verdugo, R.; Lehner, B. Intrinsic protein disorder and interaction promiscuity are widely associated with dosage sensitivity. Cell 2009, 138, 198–208.

(10) Babu, M. M. The contribution of intrinsically disordered regions to protein function, cellular complexity, and human disease. Biochemical Society Transactions 2016, 44, 1185–1200.

(11) Elbaum-Garfinkle, S. Matter over mind: Liquid phase separation and neurodegeneration. Journal of Biological Chemistry 2019, 294, 7160–7168.

(12) Uversky, V. N. Intrinsic disorder, protein–protein interactions, and disease. Advances in protein chemistry and structural biology 2018, 110, 85–121.

(13) Uversky, V. N.; Davè, V.; Iakoucheva, L. M.; Malaney, P.; Metallo, S. J.; Pathak, R. R.; Joerger, A. C. Pathological unfoldomics of uncontrolled chaos: intrinsically disordered proteins and human diseases. Chemical reviews 2014, 114, 6844–6879.

(14) Zhang, Y.; Zheng, J.; Zhang, B. Protein Language Model Identifies Disordered, Conserved Motifs Driving Phase Separation. bioRxiv 2024, 2024–12.

(15) Uversky, V. N. Intrinsically disordered proteins from A to Z. The international journal of biochemistry & cell biology 2011, 43, 1090–1103.

(16) Oldfield, C. J.; Dunker, A. K. Intrinsically disordered proteins and intrinsically disordered protein regions. Annual review of biochemistry 2014, 83, 553–584.

(17) Hansen, J. C.; Lu, X.; Ross, E. D.; Woody, R. W. Intrinsic protein disorder, amino acid composition, and histone terminal domains. Journal of Biological Chemistry 2006, 281, 1853–1856.

(18) Biesaga, M.; Frigolé-Vivas, M.; Salvatella, X. Intrinsically disordered proteins and biomolecular condensates as drug targets. Current Opinion in Chemical Biology 2021, 62, 90–100.

(19) Joshi, P.; Vendruscolo, M. Druggability of intrinsically disordered proteins. Intrinsically disordered proteins studied by NMR spectroscopy 2015, 383–400.

(20) Kosol, S.; Contreras-Martos, S.; Cedeño, C.; Tompa, P. Structural characterization of intrinsically disordered proteins by NMR spectroscopy. Molecules 2013, 18, 10802–10828.

(21) Jensen, M. R.; Zweckstetter, M.; Huang, J.-r.; Blackledge, M. Exploring free-energy landscapes of intrinsically disordered proteins at atomic resolution using NMR spectroscopy. Chemical reviews 2014, 114, 6632–6660.

(22) Gibbs, E. B.; Cook, E. C.; Showalter, S. A. Application of NMR to studies of intrinsically disordered proteins. Archives of biochemistry and biophysics 2017, 628, 57–70.

(23) Kikhney, A. G.; Svergun, D. I. A practical guide to small angle X-ray scattering (SAXS) of flexible and intrinsically disordered proteins. FEBS letters 2015, 589, 2570–2577.

(24) Sibille, N.; Bernado, P. Structural characterization of intrinsically disordered proteins by the combined use of NMR and SAXS. Biochemical society transactions 2012, 40, 955–962.

(25) Schuler, B.; Soranno, A.; Hofmann, H.; Nettels, D. Single-molecule FRET spectroscopy and the polymer physics of unfolded and intrinsically disordered proteins. Annual Review of Biophysics 2016, 45, 207–231.

(26) Regmi, R.; Srinivasan, S.; Latham, A. P.; Kukshal, V.; Cui, W.; Zhang, B.; Bose, R.; Schlau-Cohen, G. S. Phosphorylation-Dependent Conformations of the Disordered Carboxyl-Terminus Domain in the Epidermal Growth Factor Receptor. The Journal of Physical Chemistry Letters 2020, 11, 10037–10044.

(27) LeBlanc, S. J.; Kulkarni, P.; Weninger, K. R. Single molecule FRET: A powerful tool to study intrinsically disordered proteins. Biomolecules 2018, 8, 140.

(28) Tozzini, V. Coarse-grained models for proteins. Current opinion in structural biology 2005, 15, 144–150.

(29) Riniker, S.; Allison, J. R.; van Gunsteren, W. F. On developing coarse-grained models for biomolecular simulation: a review. Physical Chemistry Chemical Physics 2012, 14, 12423–12430.

(30) Dignon, G. L.; Zheng, W.; Mittal, J. Simulation methods for liquid–liquid phase separation of disordered proteins. Current opinion in chemical engineering 2019, 23, 92–98.

(31) Liu, S.; Wang, C.; Latham, A. P.; Ding, X.; Zhang, B. OpenABC enables flexible, simplified, and efficient GPU accelerated simulations of biomolecular condensates. PLoS Computational Biology 2023, 19, e1011442.

(32) Dhamankar, S.; Webb, M. A. Chemically specific coarse-graining of polymers: methods and prospects. Journal of Polymer Science 2021, 59, 2613–2643.

(33) Liu, S.; Wang, C.; Zhang, B. Toward Predictive Coarse-Grained Simulations of Biomolecular Condensates. Biochemistry 2025, 64, 1750–1761.

(34) Latham, A. P.; Zhang, B. On the Stability and Layered Organization of Protein-DNA Condensates. Biophysical Journal 2022, 121, 1727–1737.

(35) Vitalis, A.; Pappu, R. V. ABSINTH: a new continuum solvation model for simulations of polypeptides in aqueous solutions. Journal of computational chemistry 2009, 30, 673–699.

(36) Dignon, G. L.; Zheng, W.; Best, R. B.; Kim, Y. C.; Mittal, J. Relation between singlemolecule properties and phase behavior of intrinsically disordered proteins. Proceedings of the National Academy of Sciences 2018, 115, 9929–9934.

(37) Wu, H.; Wolynes, P. G.; Papoian, G. A. AWSEM-IDP: a coarse-grained force field for intrinsically disordered proteins. The Journal of Physical Chemistry B 2018, 122, 11115–11125.

(38) Joseph, J. A.; Reinhardt, A.; Aguirre, A.; Chew, P. Y.; Russell, K. O.; Espinosa, J. R.; Garaizar, A.; Collepardo-Guevara, R. Physics-driven coarse-grained model for biomolecular phase separation with near-quantitative accuracy. Nature Computational Science 2021, 1, 732–743.

(39) Latham, A. P.; Zhang, B. Consistent force field captures homologue-resolved hp1 phase separation. Journal of chemical theory and computation 2021, 17, 3134–3144.

(40) Regy, R. M.; Thompson, J.; Kim, Y. C.; Mittal, J. Improved coarse-grained model for studying sequence dependent phase separation of disordered proteins. Protein Science 2021, 30, 1371–1379.

(41) Dannenhoffer-Lafage, T.; Best, R. B. A data-driven hydrophobicity scale for predicting liquid–liquid phase separation of proteins. The Journal of Physical Chemistry B 2021, 125, 4046–4056.

(42) Tesei, G.; Schulze, T. K.; Crehuet, R.; Lindorff-Larsen, K. Accurate model of liquid–liquid phase behavior of intrinsically disordered proteins from optimization of single-chain properties. Proceedings of the National Academy of Sciences 2021, 118, e2111696118.

(43) Zhang, Y.; Liu, X.; Chen, J. Toward accurate coarse-grained simulations of disordered proteins and their dynamic interactions. Journal of chemical information and modeling 2022, 62, 4523–4536.

(44) Zhang, Y.; Li, S.; Gong, X.; Chen, J. Toward accurate simulation of coupling between protein secondary structure and phase separation. Journal of the American Chemical Society 2023, 146, 342–357.

(45) Thomasen, F. E.; Pesce, F.; Roesgaard, M. A.; Tesei, G.; Lindorff-Larsen, K. Improving Martini 3 for disordered and multidomain proteins. Journal of chemical theory and computation 2022, 18, 2033–2041.

(46) Tesei, G.; Lindorff-Larsen, K. Improved predictions of phase behaviour of intrinsically disordered proteins by tuning the interaction range. Open Research Europe 2023, 2, 94.

(47) Cao, F.; von Bülow, S.; Tesei, G.; Lindorff-Larsen, K. A coarse-grained model for disordered and multi-domain proteins. Protein Science 2024, 33, e5172.

(48) Souza, P. C. et al. Martini 3: A General Purpose Force Field for Coarse-Grained Molecular Dynamics. Nature Methods 2021, 18, 382–388.

(49) Ding, X.; Zhang, B. Contrastive Learning of Coarse-Grained Force Fields. Journal of Chemical Theory and Computation 2022, 18, 6334–6344.

(50) Ding, X. Optimizing force fields with experimental data using ensemble reweighting and potential contrasting. The Journal of Physical Chemistry B 2024, 128, 6760–6769.

(51) R. Tejedor, A.; Aguirre Gonzalez, A.; Maristany, M. J.; Chew, P. Y.; Russell, K.; Ramirez, J.; Espinosa, J. R.; Collepardo-Guevara, R. Chemically Informed Coarse-Graining of Electrostatic Forces in Charge-Rich Biomolecular Condensates. ACS Central Science 2025, 11, 302–321.

(52) Airas, J.; Ding, X.; Zhang, B. Transferable Implicit Solvation via Contrastive Learning of Graph Neural Networks. ACS Cent. Sci. 2023, 9, 2286–2297.

(53) Lotthammer, J. M.; Ginell, G. M.; Griffith, D.; Emenecker, R.; Holehouse, A. S. Direct prediction of intrinsically disordered protein conformational properties from sequence. Biophysical Journal 2024, 123, 43a.

(54) Chakraborty, D.; Mondal, B.; Thirumalai, D. Brewing coffee: A sequence-specific coarse-grained energy function for simulations of DNA-protein complexes. Journal of Chemical Theory and Computation 2024, 20, 1398–1413.

(55) Jumper, J.; Evans, R.; Pritzel, A.; Green, T.; Figurnov, M.; Ronneberger, O.; Tunyasuvunakool, K.; Bates, R.; Žídek, A.; Potapenko, A.; others Highly accurate protein structure prediction with AlphaFold. nature 2021, 596, 583–589.

(56) Baek, M.; DiMaio, F.; Anishchenko, I.; Dauparas, J.; Ovchinnikov, S.; Lee, G. R.; Wang, J.; Cong, Q.; Kinch, L. N.; Schaeffer, R. D.; others Accurate prediction of protein structures and interactions using a three-track neural network. Science 2021, 373, 871–876.

(57) Janson, G.; Feig, M. Transferable deep generative modeling of intrinsically disordered protein conformations. PLOS Computational Biology 2024, 20, e1012144.

(58) Janson, G.; Valdes-Garcia, G.; Heo, L.; Feig, M. Direct generation of protein conformational ensembles via machine learning. Nature Communications 2023, 14, 774.

(59) Zhu, J.; Li, Z.; Zheng, Z.; Zhang, B.; Zhong, B.; Bai, J.; Hong, X.; Wang, T.; Wei, T.; Yang, J.; others Precise generation of conformational ensembles for intrinsically disordered proteins via fine-tuned diffusion models. bioRxiv 2024, 2024–05.

(60) Novak, B.; Lotthammer, J. M.; Emenecker, R. J.; Holehouse, A. S. Accurate predictions of conformational ensembles of disordered proteins with STARLING. bioRxiv 2025, 2025–02.

(61) Mao, A. H.; Crick, S. L.; Vitalis, A.; Chicoine, C. L.; Pappu, R. V. Net charge per residue modulates conformational ensembles of intrinsically disordered proteins. Proceedings of the National Academy of Sciences 2010, 107, 8183–8188.

(62) Das, R. K.; Pappu, R. V. Conformations of intrinsically disordered proteins are influenced by linear sequence distributions of oppositely charged residues. Proceedings of the National Academy of Sciences 2013, 110, 13392–13397.

(63) Müller-Späth, S.; Soranno, A.; Hirschfeld, V.; Hofmann, H.; Rüegger, S.; Reymond, L.; Nettels, D.; Schuler, B. Charge interactions can dominate the dimensions of intrinsically disordered proteins. Proceedings of the National Academy of Sciences 2010, 107, 14609–14614.

(64) Das, R. K.; Ruff, K. M.; Pappu, R. V. Relating sequence encoded information to form and function of intrinsically disordered proteins. Current opinion in structural biology 2015, 32, 102–112.

(65) Zheng, W.; Dignon, G.; Brown, M.; Kim, Y. C.; Mittal, J. Hydropathy patterning complements charge patterning to describe conformational preferences of disordered proteins. The journal of physical chemistry letters 2020, 11, 3408–3415.

(66) Holla, A.; Martin, E. W.; Dannenhoffer-Lafage, T.; Ruff, K. M.; König, S. L.; Nüesch, M. F.; Chowdhury, A.; Louis, J. M.; Soranno, A.; Nettels, D.; others Identifying Sequence Effects on Chain Dimensions of Disordered Proteins by Integrating Experiments and Simulations. JACS Au 2024, 4, 4729–4743.

(67) Baul, U.; Chakraborty, D.; Mugnai, M. L.; Straub, J. E.; Thirumalai, D. Sequence effects on size, shape, and structural heterogeneity in intrinsically disordered proteins. The Journal of Physical Chemistry B 2019, 123, 3462–3474.

(68) Zheng, W.; Dignon, G. L.; Jovic, N.; Xu, X.; Regy, R. M.; Fawzi, N. L.; Kim, Y. C.; Best, R. B.; Mittal, J. Molecular details of protein condensates probed by microsecond long atomistic simulations. The Journal of Physical Chemistry B 2020, 124, 11671–11679.

(69) Galvanetto, N.; Ivanović, M. T.; Chowdhury, A.; Sottini, A.; Nüesch, M. F.; Nettels, D.; Best, R. B.; Schuler, B. Extreme dynamics in a biomolecular condensate. Nature 2023, 619, 876–883.

(70) Rekhi, S.; Garcia, C. G.; Barai, M.; Rizuan, A.; Schuster, B. S.; Kiick, K. L.; Mittal, J. Expanding the molecular language of protein liquid–liquid phase separation. Nature Chemistry 2024, 16, 1113–1124.

(71) Wang, C.; Kilgore, H. R.; Latham, A. P.; Zhang, B. Nonspecific Yet Selective Interactions Contribute to Small Molecule Condensate Binding. J. Chem. Theory Comput. 2024, 20, 10247–10258.

(72) Latham, A. P.; Zhu, L.; Sharon, D. A.; Ye, S.; Willard, A. P.; Zhang, X.; Zhang, B. Microphase Separation Produces Interfacial Environment within Diblock Biomolecular Condensates. eLife 2024, 12 .

(73) Zhou, L.; Zhu, L.; Wang, C.; Xu, T.; Wang, J.; Zhang, B.; Zhang, X.; Wang, H. Multiphasic Condensates Formed with Mono-Component of Tetrapeptides via Phase Separation. Nat Commun 2025, 16, 2706.

(74) McInnes, L.; Healy, J.; Melville, J. Umap: Uniform manifold approximation and projection for dimension reduction. arXiv preprint arXiv:1802.03426 2018,

(75) Van Der Lee, R. et al. Classification of Intrinsically Disordered Regions and Proteins. Chemical Reviews 2014, 114, 6589–6631.

(76) Abyzov, A.; Blackledge, M.; Zweckstetter, M. Conformational Dynamics of Intrinsically Disordered Proteins Regulate Biomolecular Condensate Chemistry. Chemical Reviews 2022, 122, 6719–6748.

(77) Zarin, T.; Strome, B.; Nguyen Ba, A. N.; Alberti, S.; Forman-Kay, J. D.; Moses, A. M. Proteome-wide signatures of function in highly diverged intrinsically disordered regions. Elife 2019, 8, e46883.

(78) Mirdita, M.; Schütze, K.; Moriwaki, Y.; Heo, L.; Ovchinnikov, S.; Steinegger, M. ColabFold: making protein folding accessible to all. Nature methods 2022, 19, 679–682.

(79) Wilson, C. J.; Choy, W.-Y.; Karttunen, M. AlphaFold2: a role for disordered protein/region prediction? International Journal of Molecular Sciences 2022, 23, 4591.

(80) Robustelli, P.; Piana, S.; Shaw, D. E. Developing a molecular dynamics force field for both folded and disordered protein states. Proceedings of the National Academy of Sciences 2018, 115, E4758–E4766.

(81) Chen, J.; Liu, X.; Chen, J. Targeting intrinsically disordered proteins through dynamic interactions. Biomolecules 2020, 10, 743.

(82) Liu, X.; Chen, J. Residual structures and transient long-range interactions of p53 transactivation domain: Assessment of explicit solvent protein force fields. Journal of chemical theory and computation 2019, 15, 4708–4720.

(83) Mukherjee, S.; Schäfer, L. V. Thermodynamic forces from protein and water govern condensate formation of an intrinsically disordered protein domain. Nature Communications 2023, 14, 5892.

(84) Kyte, J.; Doolittle, R. F. A simple method for displaying the hydropathic character of a protein. Journal of molecular biology 1982, 157, 105–132.

(85) Uversky, V. N.; Gillespie, J. R.; Fink, A. L. Why are “natively unfolded” proteins unstructured under physiologic conditions? Proteins: structure, function, and bioinformatics 2000, 41, 415–427.

(86) Le Guillou, J.; Zinn-Justin, J. Critical exponents for the n-vector model in three dimensions from field theory. Physical Review Letters 1977, 39, 95.

(87) Gates, Z. P.; Baxa, M. C.; Yu, W.; Riback, J. A.; Li, H.; Roux, B.; Kent, S. B.; Sosnick, T. R. Perplexing cooperative folding and stability of a low-sequence complexity, polyproline 2 protein lacking a hydrophobic core. Proceedings of the National Academy of Sciences 2017, 114, 2241–2246.

(88) Shrestha, U. R.; Smith, J. C.; Petridis, L. Full structural ensembles of intrinsically disordered proteins from unbiased molecular dynamics simulations. Communications biology 2021, 4, 243.

(89) Burke, K. A.; Janke, A. M.; Rhine, C. L.; Fawzi, N. L. Residue-by-residue view of in vitro FUS granules that bind the C-terminal domain of RNA polymerase II. Molecular cell 2015, 60, 231–241.

(90) Grosberg, A. Statistical Physics of Macromolecules; AIP Press: New York, 1994.

(91) Tesei, G.; Trolle, A. I.; Jonsson, N.; Betz, J.; Knudsen, F. E.; Pesce, F.; Johansson, K. E.; Lindorff-Larsen, K. Conformational ensembles of the human intrinsically disordered proteome. Nature 2024, 626, 897–904.

(92) Abraham, M. J.; Murtola, T.; Schulz, R.; Páll, S.; Smith, J. C.; Hess, B.; Lindahl, E. GROMACS: High performance molecular simulations through multi-level parallelism from laptops to supercomputers. SoftwareX 2015, 1, 19–25.

(93) Eastman, P.; Swails, J.; Chodera, J. D.; McGibbon, R. T.; Zhao, Y.; Beauchamp, K. A.; Wang, L.-P.; Simmonett, A. C.; Harrigan, M. P.; Stern, C. D.; others OpenMM 7: Rapid development of high performance algorithms for molecular dynamics. PLoS computational biology 2017, 13, e1005659.

(94) Hopkins, C. W.; Le Grand, S.; Walker, R. C.; Roitberg, A. E. Long-time-step molecular dynamics through hydrogen mass repartitioning. Journal of chemical theory and computation 2015, 11, 1864–1874.

(95) Zhang, Z.; Liu, X.; Yan, K.; Tuckerman, M. E.; Liu, J. Unified efficient thermostat scheme for the canonical ensemble with holonomic or isokinetic constraints via molecular dynamics. The Journal of Physical Chemistry A 2019, 123, 6056–6079.

(96) Stefko, M.; Douglass, K.; Manley, S. PolymerCpp (0.1.3). 2020; 10.5281/zenodo.3928659.

(97) McGibbon, R. T.; Beauchamp, K. A.; Harrigan, M. P.; Klein, C.; Swails, J. M.; Hernández, C. X.; Schwantes, C. R.; Wang, L.-P.; Lane, T. J.; Pande, V. S. MDTraj: a modern open library for the analysis of molecular dynamics trajectories. Biophysical journal 2015, 109, 1528–1532.

(98) Lalmansingh, J. M.; Keeley, A. T.; Ruff, K. M.; Pappu, R. V.; Holehouse, A. S. SOURSOP: A Python package for the analysis of simulations of intrinsically disordered proteins. Journal of Chemical Theory and Computation 2023, 19, 5609–5620.

