## Supporting Information for "Sequence-Dependent Conformational Landscapes of Intrinsically Disordered Proteins"

### Supporting Information: Sequence-Dependent Conformational Landscapes of Intrinsically Disordered Proteins

### Contents

**IDP sequences** **S-3**

**Supplemental figures** **S-5**

**References** **S-23**

### IDP sequences

IDP 1: ELDPESEVQAVGTGTKTDSDSTKATSAETAGKTSSAILED

IDP 2: QSISSRVSYNLDICFLQQAISSINDFKGLENKYCVLGREA

IDP 3: ITHETRTTPSVNTINKSNPQGPSDNATSIKKTLNRVPKWM

IDP 4: DWALEVENEEIEQTQPTPLEGEDEEEGLDSEFKKKITLQD

IDP 5: DCNFLDDSTVTNISANHSITSLVNEWAEYDLLSDTDLSSN

IDP 6: FLEECREKEEFVNESTDDEESKFDIADYKDEMVSDEHLWN

IDP 7: EESLIESEEATREEVDDFVGSDDAVALGDMRPQQDYGRIL

IDP 8: RFKNNNHTISVFDHDLSTVTEPIQIMDTLEATTAPGNDER

IDP 9: DYEALKAQQDKGKKNGFGKGQVLDPSVLGQGSMSTRIDYA

IDP 10: IDQVSNDSNDKKGASDMPTNTENVLRNRSEGILRQPEIER

IDP 11: MPVLKSDNFDPLEEAYEGGTIQNYNDEHHLHKSWANVIPD

IDP 12: TNSGSACGSVLSVNSLSNKRSRADDYDCMDEDKENQPHNP

IDP 13: IFDTSEKQCSVIKQNYGIAYGSHGCVIDITGDLCITNACM

IDP 14: VPHLVNQTKNETEEDASETGTIDSHLDRKKKKEGAFKILL

IDP 16: VLYDEGYEDKKSLHFADKKSIRTGYASNTYADDEDDDFFD

IDP 17: MLSTRTKGAICLGRKTHIGAKTLTLGQNYTTSSHAHTTTN

IDP 18: NKFRDYKESDHYAGGAGGYNNASSMDTTHNDKSTYDSDKH

IDP 19: VKKHASPDFIFVLSERIKLPDIDSEDKPELPRTNSYKFMM

IDP 20: PYTQTYTNPSHCQLCVPSATSRSASSASLSQQLQNSPTGC

IDP 21: LWLREHSKEAPPGAPGAPSSPSELECSANPQLNDKDPQYV

IDP 22: STESARSSHDHLNNDNRMFTAETSGITDPATQANMGNQIT

IDP 23: MNSTLGTSTIPRGLIHRPSAGRTTRPCDNCTCSPGLLSRQ

IDP 24: DAAAAAGLPTTSGSGGGNAGPGAGDEEESGKKTEKDSALM

IDP 25: LLVPENSRPPVQALPKEYQVRPRTTYEDGPGTPEWKRARL

IDP 26: MFNMNLLSTPSSEEGSPQNRSSSMSSVEGKKDRDTFTNLQ

IDP 30: SNAVATDTGISRNTGSNTTKEQKFSAIDATDSQNDGSYGG

IDP 31: TMTSPSGSGRVKNIYNSSNGSPSPSGWDSPSFRNRYNFDD

IDP 32: FYSPSTDTYLNNGGMQPSNGSVYSANTYRGQVNGAPLPYQ

IDP 33: NTPIASGILQPAAFDYFSRPISTQDIISNNCGNTLSRGPL

IDP 34: AGGANVSPSSGSTPHILPSLPTSTSNASSGPPYGYPQPAH

IDP 35: MSYNPYAYAPDNGQPVTDRFAQHPQAQQQTNELPKQPSPG

IDP 36: MLSEELYDSGSTATSTNSLFSLNKRGSYTSVVDCYLDEAE

IDP 37: TCVPTSTGINLGSSSSNSGSGRGDSTTATSSAAAAADAIQ

IDP 38: MNIFSQVGGLSPNYDKNFISNGDVHGRETDDGEFDDSVNM

IDP 39: KSTEHQQKGSAAVNSYAEGQDIGNGMGFSLNTRSFSEPAW

IDP 40: SYVPLLSSVVPGKRHSDVLCPSLSESAISDDDEHEPIVRC

IDP 41: MSETTNTLRRRTNEEWSATAGIAEQHENQPSVSTDKKDQS

IDP 42: DDNPKDPKSYDKRDGSNVVDTSKPGDGNQGNDMDWLFRNA

IDP 44: PRPKRQLEEDTNKKQKKKKKFPQKAAVVESDAKSSEMGEG

IDP 45: EHGFNYDRTNSTDSSSVNLTSVDQRVSSRQKSGSTITLNT

IDP 46: ICSSEFKDNYYQKVESPTRTPNDWKKNNLLSKNKNTENNK

IDP 47: DLNSKRYSNIPSSKPAGEALSPVRSHNSGEYRRADMMTGK

IDP 48: MVSISILKGKKKGTERPIEVTHHSYAGGRHEKTKRGTAGV

IDP 49: MFNRSRTNKKLLPNFKEKDEENKIGKVRKDSLDASKPLRL

IDP 51: MSASEAGVTEQVKKLSVKDSSNDAVKPNKKENKKSKQQSL

IDP 52: MKAFTSLLCGLGLSTTLAKAISLQRPLGLDKDVLLQAAEK

IDP 53: MNWLLRRGTCWTLAPAWLRCRCPSSSRRLYSLAHEVDTSK

#### Supplemental figures

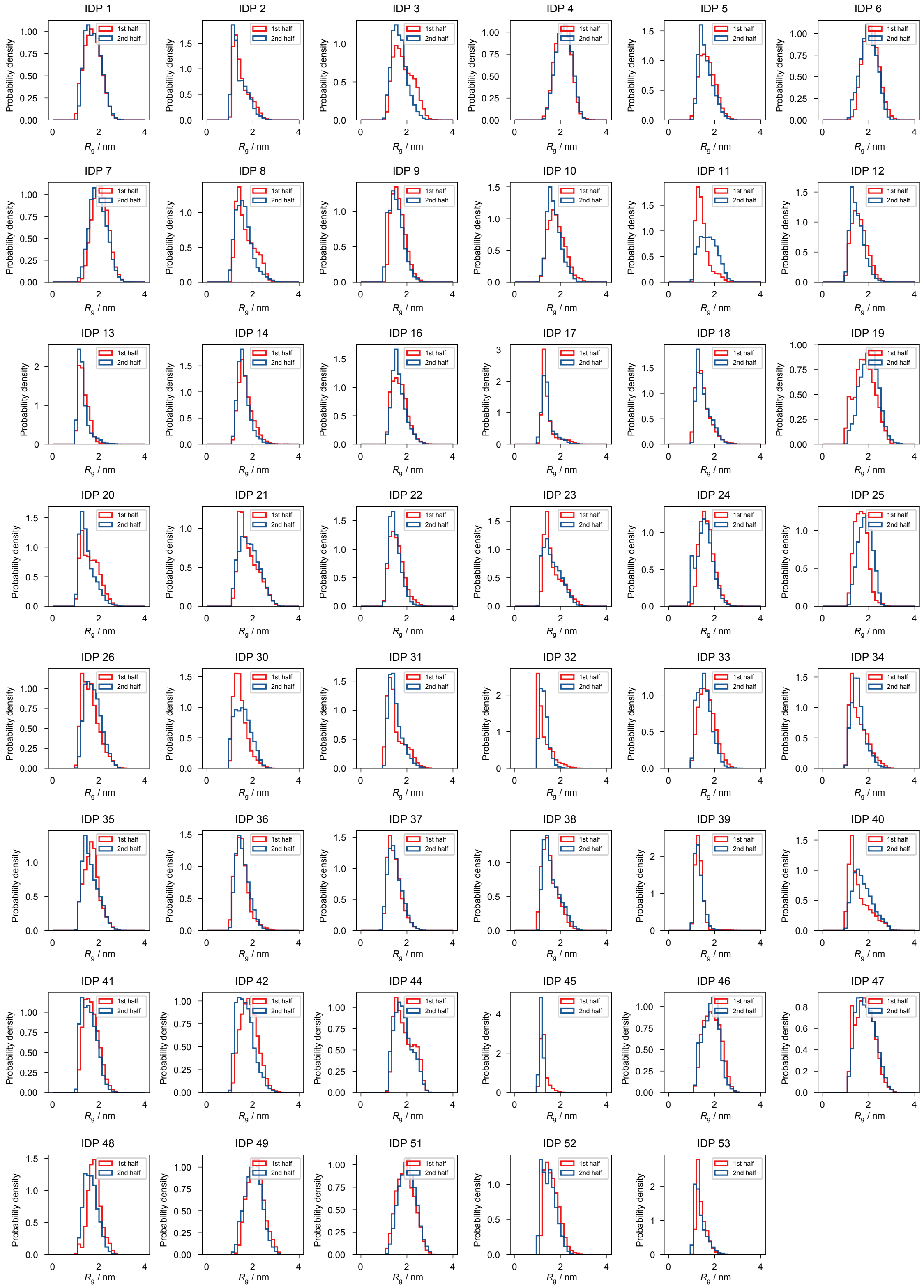


Figure S1: **Probability distributions of the radius of gyration for each IDP ensemble, computed separately for the first and second halves of the simulation.** The substantial overlap between distributions indicates that the simulations are well-converged.

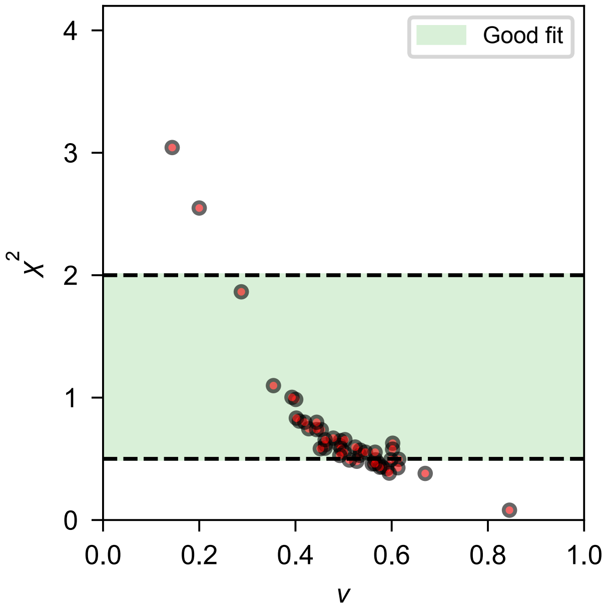


Figure S2: **Fitted Flory–Huggins scaling exponents versus reduced $\chi^2$ values.** Each point represents a single IDP ensemble, with fitted values obtained from a log–log linear regression of the form $y = \nu x + \log(A)$, where $x = \log(|i - j|)$ is the logarithm of the sequence separation between residues and $y = \log(\langle r_{ij}^2 \rangle^{1/2})$ is the logarithm of the ensemble-averaged root-mean-square spatial distance. Here, $\nu$ denotes the Flory–Huggins scaling exponent and $A$ is the prefactor. The reduced $\chi^2$ statistic for each fit was calculated as $\chi^2_{\text{reduced}} = \frac{1}{n-p} \sum_{i=1}^{n} \frac{(y_i - y_{\text{model}}(x_i))^2}{\sigma_i^2}$, where $n$ is the number of data points, $p = 2$ is the number of fitted parameters, and $\sigma_i^2$ is the estimated variance in $y_i$. The model function $y_{\text{model}}(x_i)$ corresponds to the fitted scaling relation.[S1] See the `SOURSOP` package[S2] for implementation details. The green shaded region indicates the conventional range for acceptable fits ($\chi^2_{\text{reduced}} \sim 1$). Most fitted exponents fall between 0.5 and 0.6, consistent with previously reported values for IDPs (mean $\nu \approx 0.588$). Fits in this range are generally associated with $\chi^2_{\text{reduced}}$ values near one, indicating good agreement with the scaling model. In contrast, extreme exponent values correspond to substantially elevated or suppressed $\chi^2_{\text{reduced}}$, indicating poor fit quality.

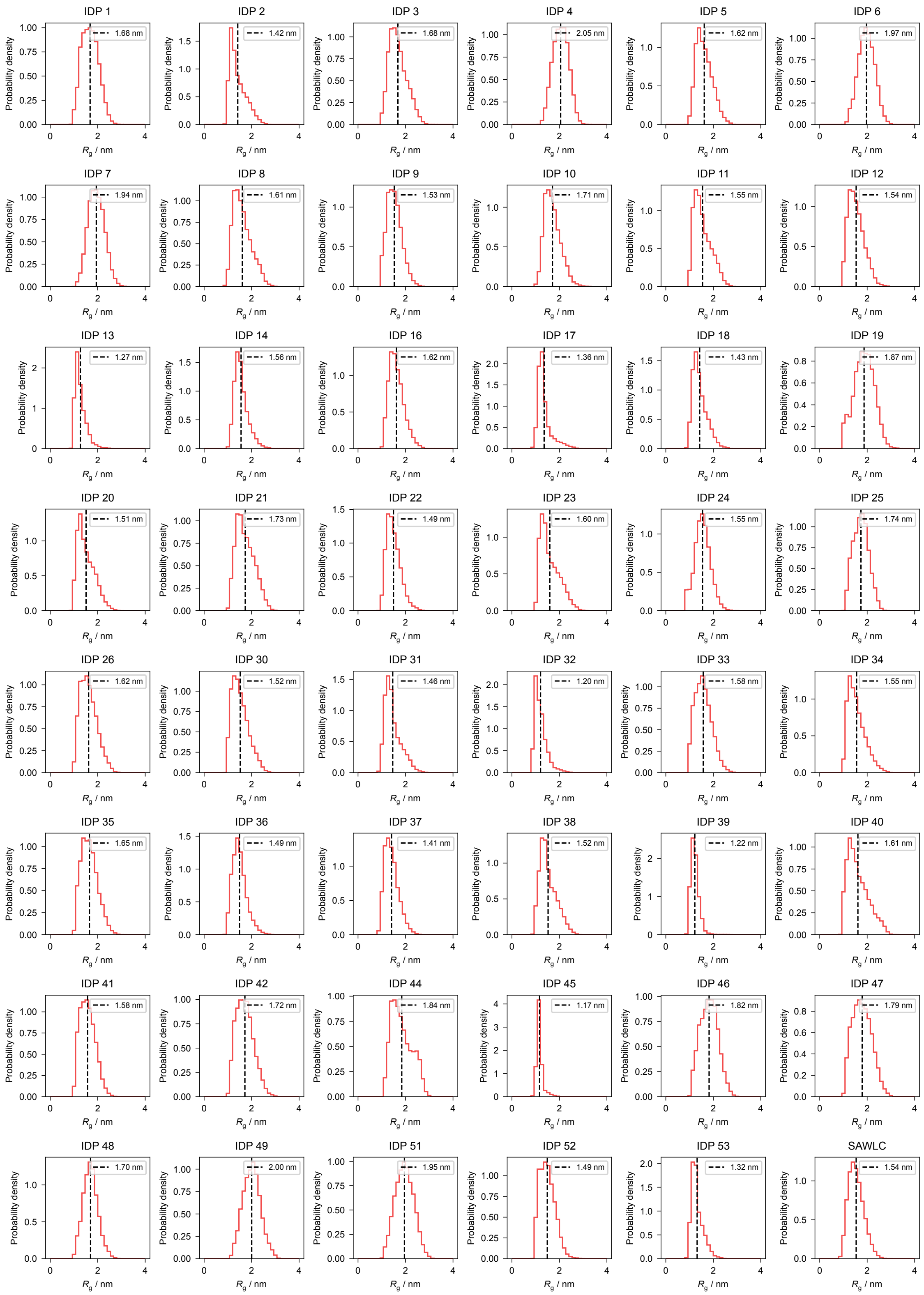


Figure S3: **Probability distributions of radius of gyration for individual IDP ensembles.** Each panel represents one IDP, and the dashed vertical line indicates the ensemble-averaged $R_{\rm g}$ value.

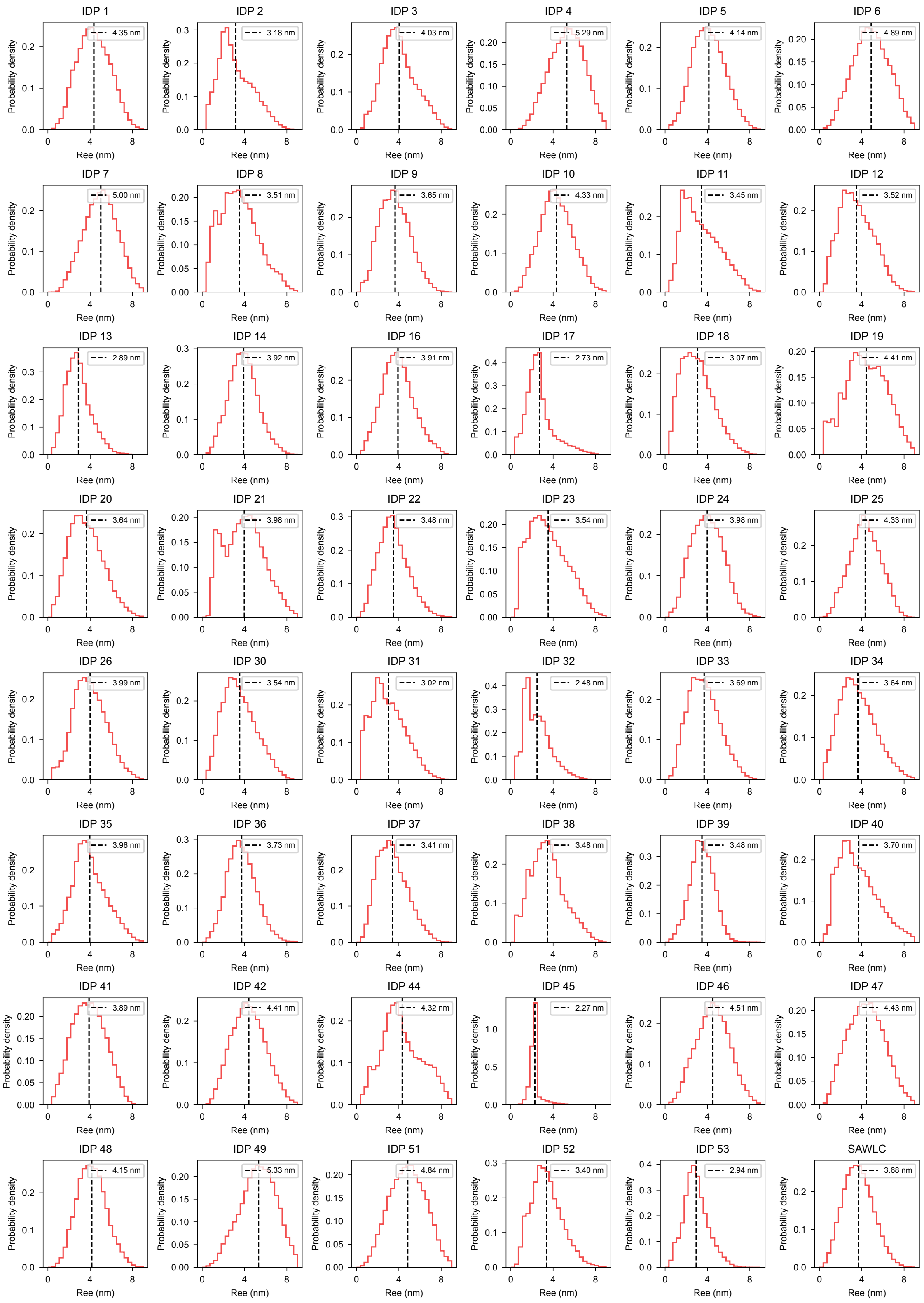


Figure S4: **Probability distributions of end-to-end distance for individual IDP ensembles.** Each panel represents one IDP, and the dashed vertical line indicates the ensemble-averaged $R_{\mathrm{ee}}$ value.

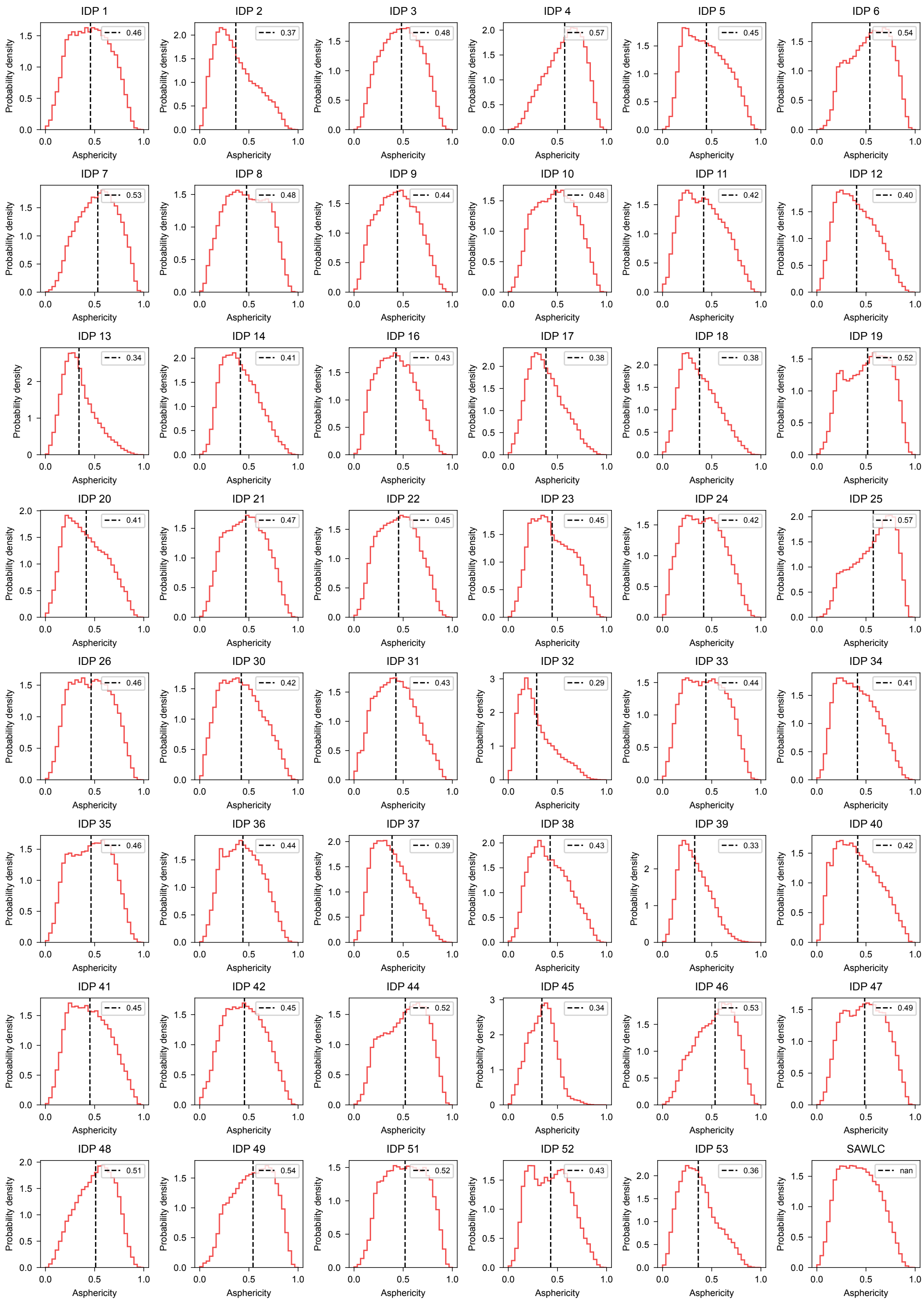


Figure S5: **Probability distributions of asphericity for individual IDP ensembles.** Each panel represents one IDP, and the dashed vertical line indicates the ensemble-averaged asphericity value.

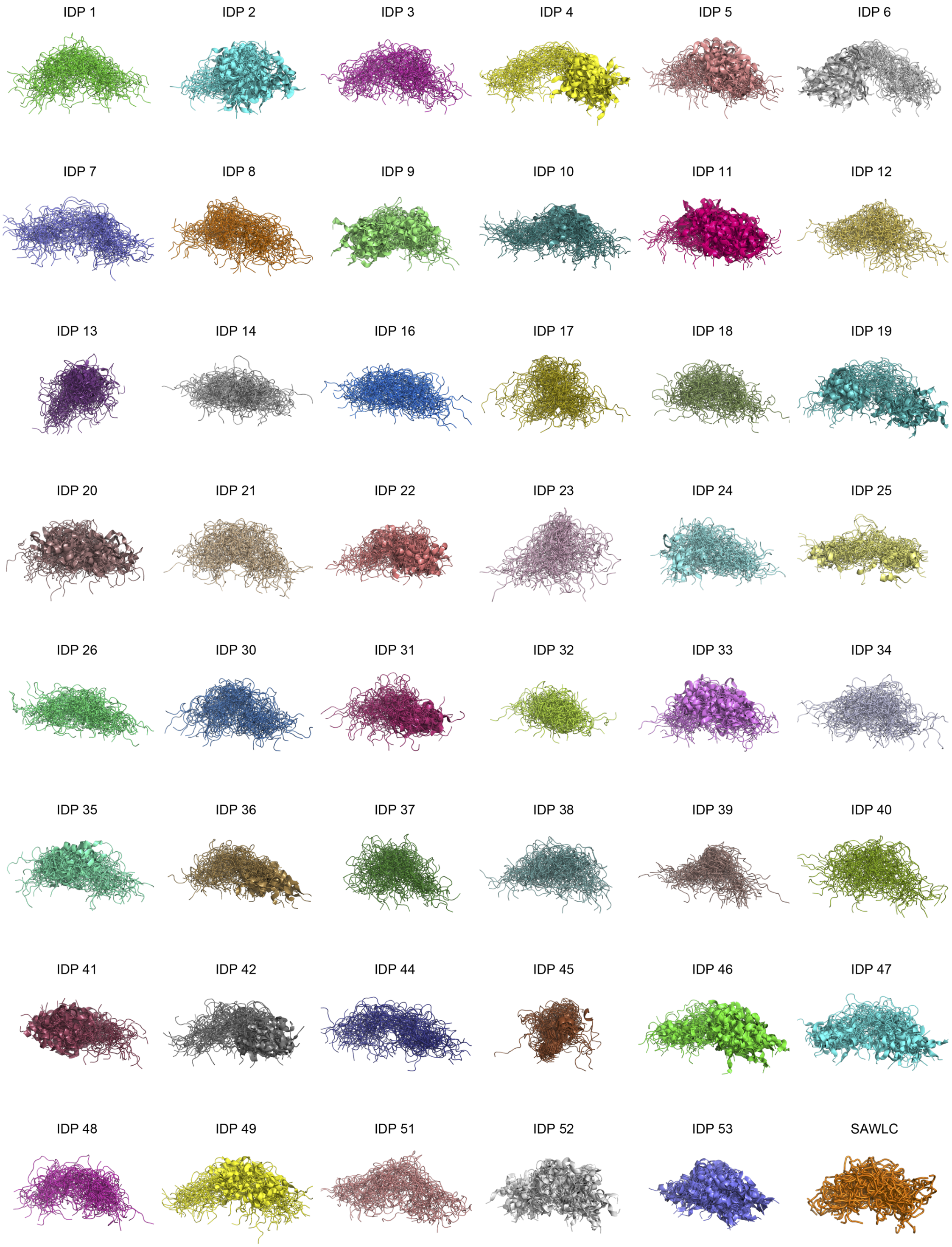


Figure S6: **Representative conformations from individual IDP ensembles.** For each ensemble, 53 conformations were uniformly sampled from a 105,000-frame trajectory to visualize the diversity of its conformational landscape.

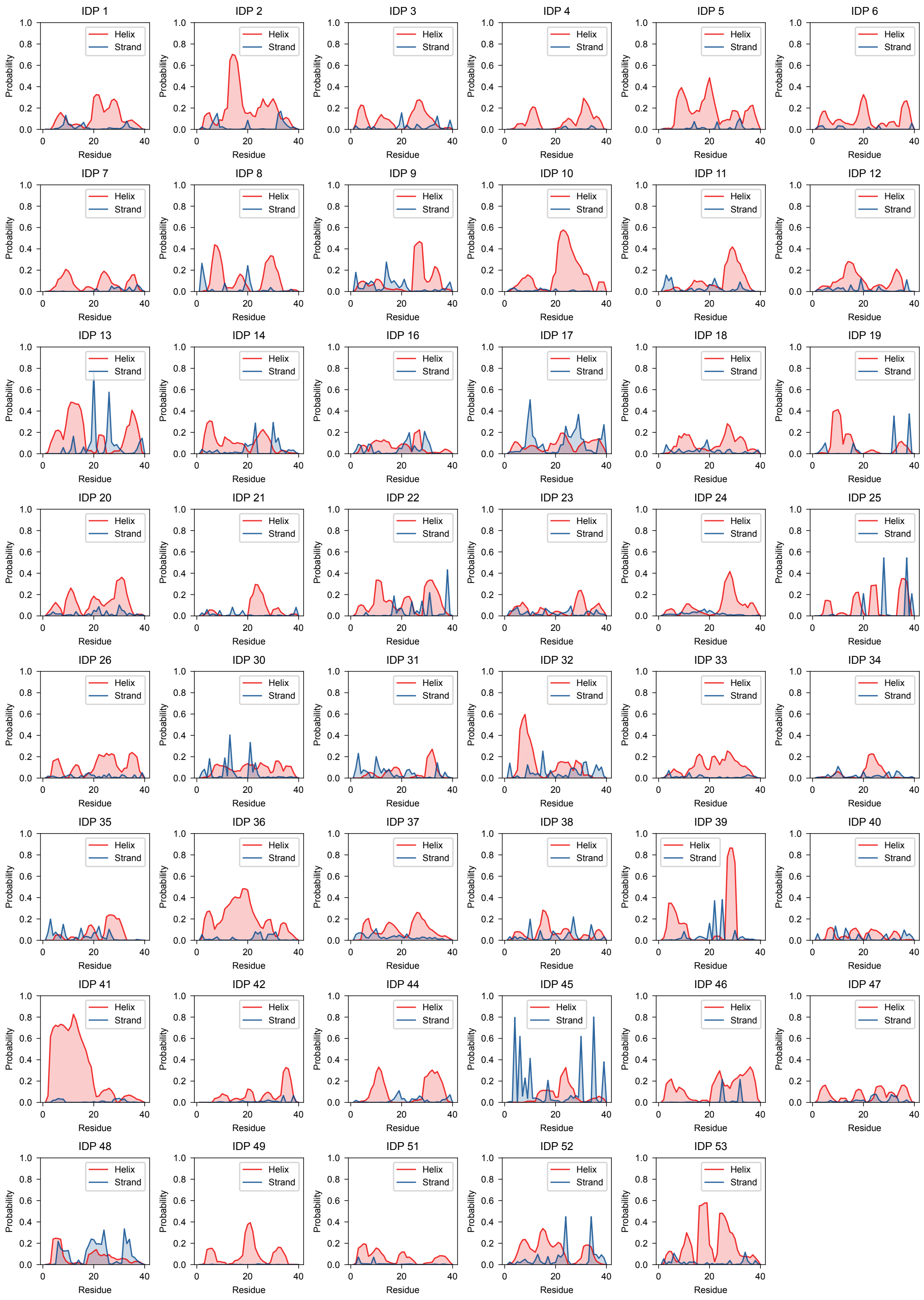


Figure S7: **Ensemble-averaged residue-wise probabilities of helix and beta-strand formation for individual IDP ensembles.** Each panel represents one IDP, with secondary structure propensities computed over the entire simulation trajectory.

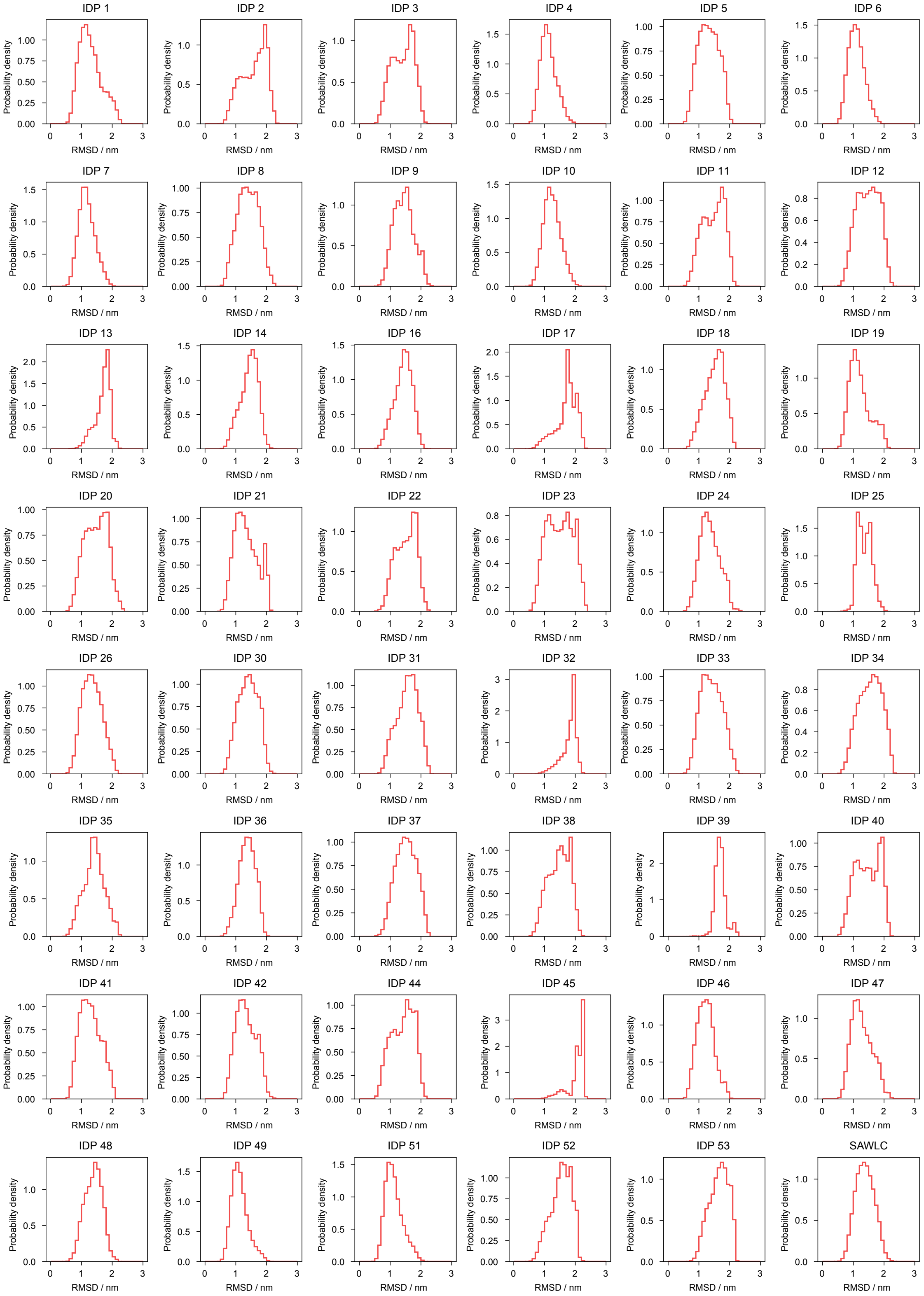


Figure S8: **Probability distributions of RMSD for individual IDP ensembles.** Each panel represents one IDP, and the RMSD values were computed using the initial structure of IDP 1 as the reference.

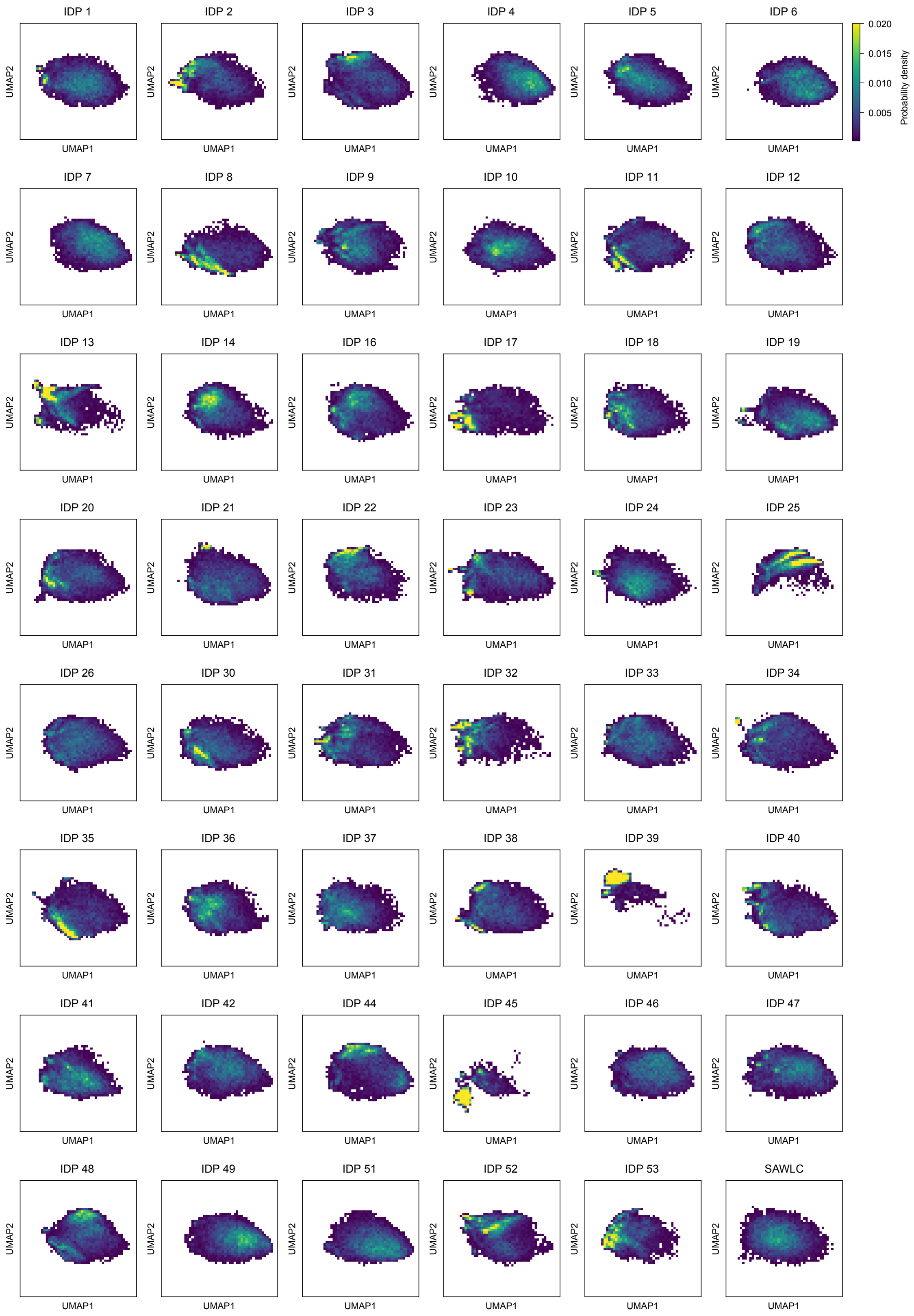


Figure S9: **Probability distributions of UMAP embeddings for individual IDP ensembles.** Each panel represents one IDP.

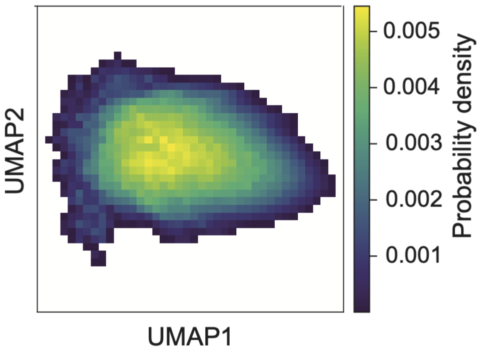


Figure S10: **Probability distributions of UMAP embeddings computed from the combined conformational ensembles of all IDPs.**

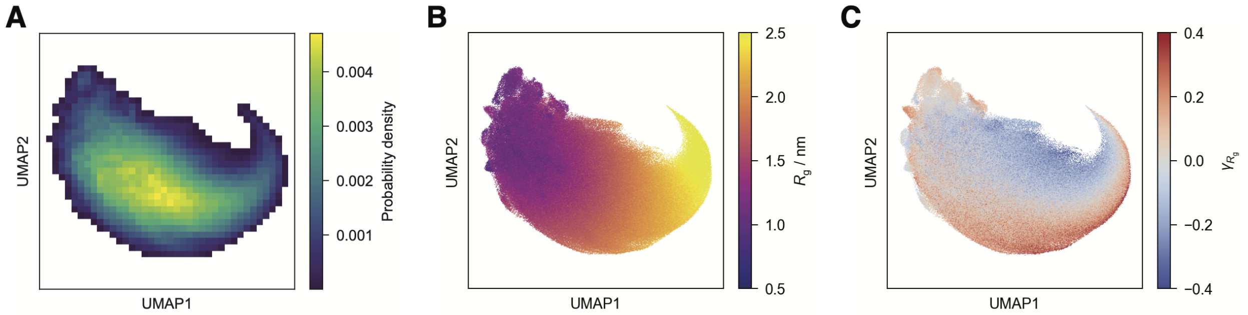


Figure S11: **UMAP embeddings computed using RMSD-based descriptors exhibit similar properties to those based on pairwise distances.** (A) Probability density distributions of UMAP embeddings derived from the combined conformational ensembles of all IDPs. (B) UMAP embedding of all conformational ensembles, with each point colored by the corresponding radius of gyration ($R_\mathrm{g}$). (C) UMAP embedding of all conformational ensembles, with each point colored by local compactness asymmetry ($\gamma_{R_\mathrm{g}}$).

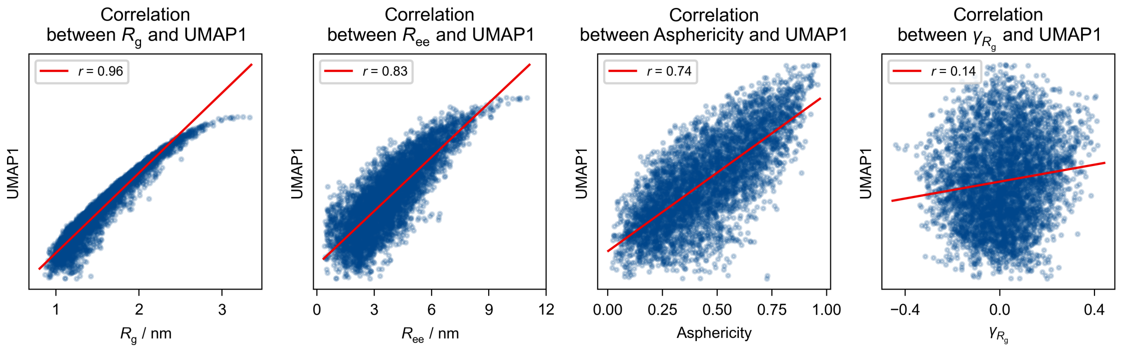


Figure S12: **Correlation between UMAP1 and various conformational descriptors across all IDP ensembles.**

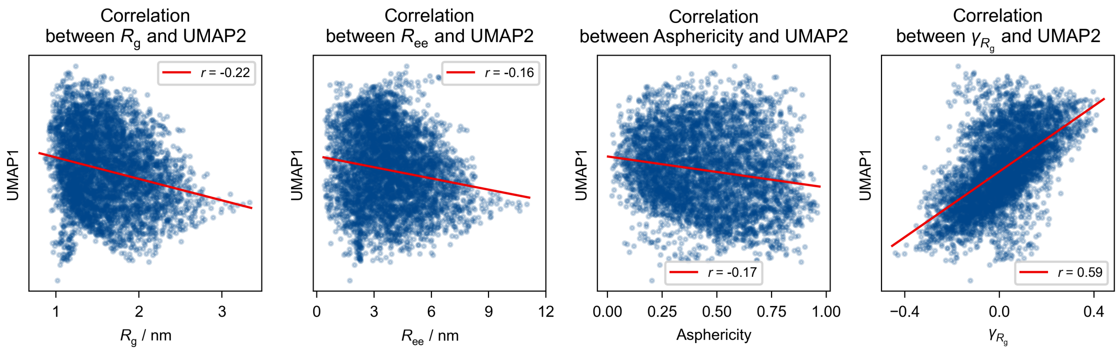


Figure S13: **Correlation between UMAP2 and various conformational descriptors across all IDP ensembles.**

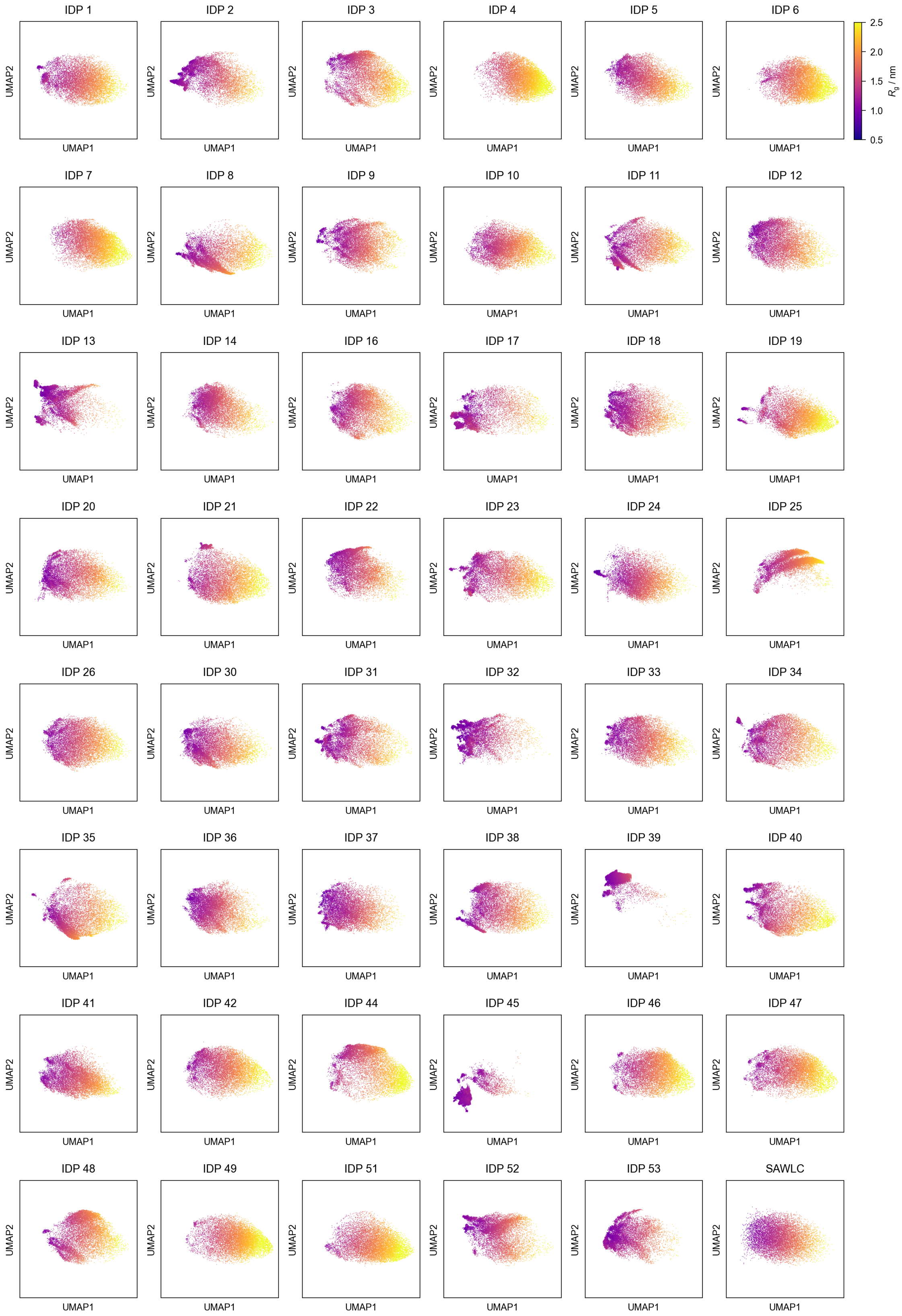


Figure S14: **Scatter plot of UMAP embeddings for individual IDP ensembles, with each point colored by the corresponding radius of gyration ($R_g$).** Each panel represents one IDP.

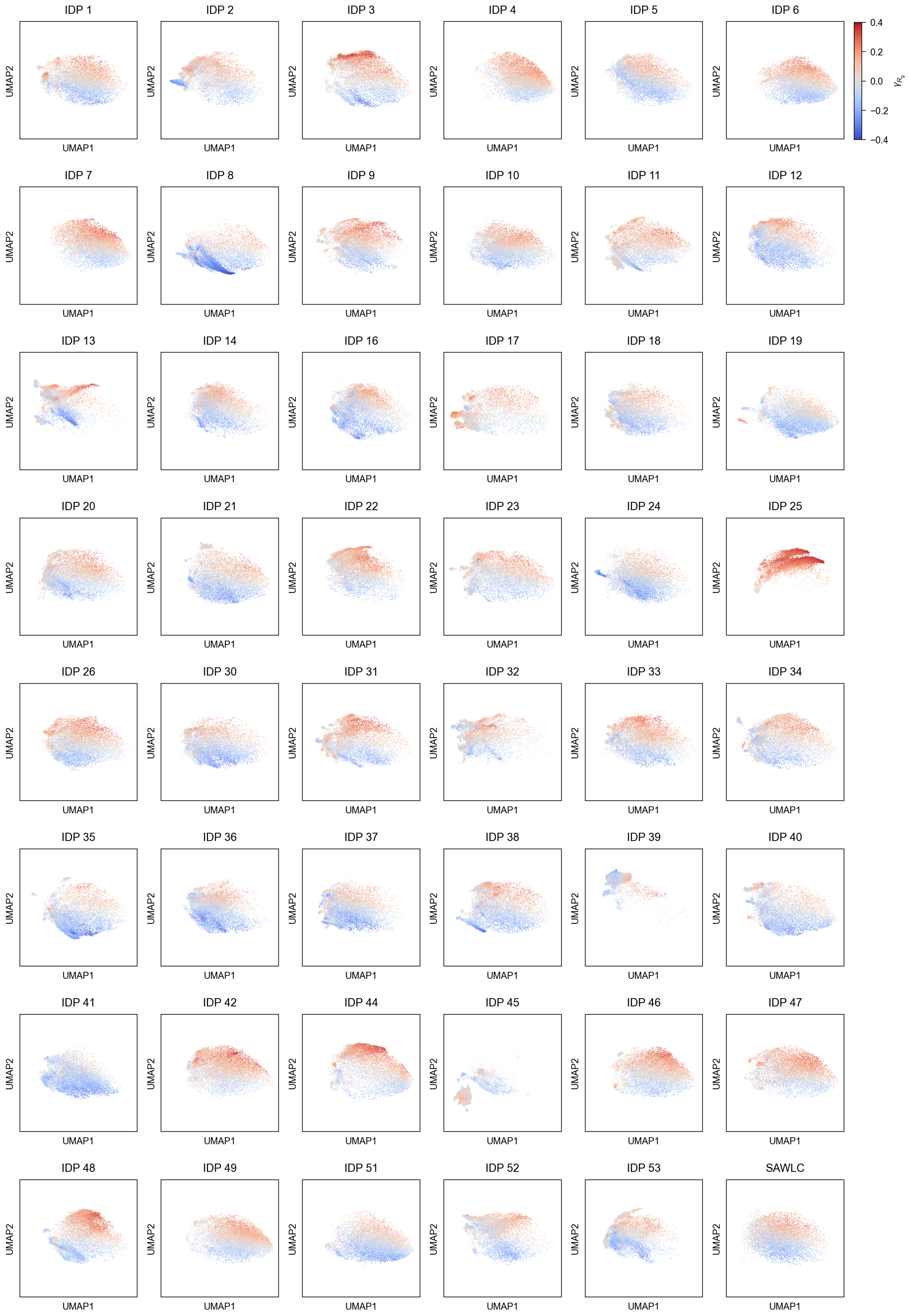


Figure S15: **Scatter plot of UMAP embeddings for individual IDP ensembles, with each point colored by local compactness asymmetry ($\gamma_{R_g}$).** Each panel represents one IDP.

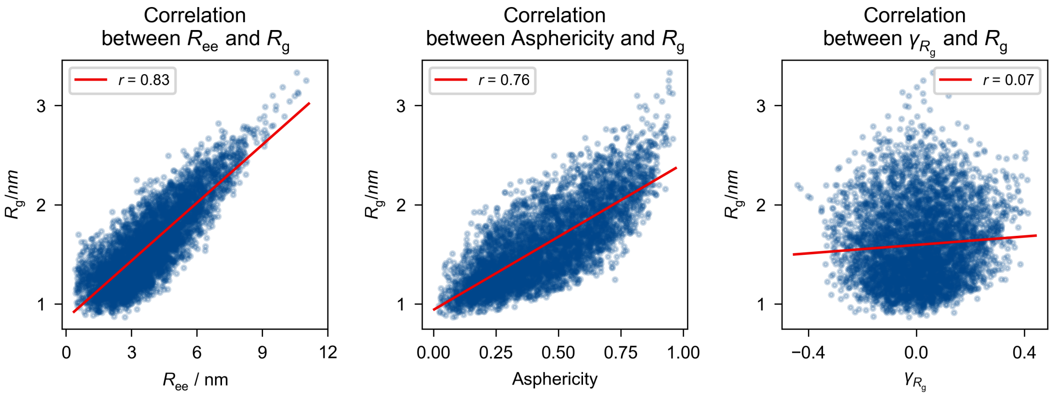


Figure S16: **Correlation between $R_g$ and various conformational descriptors across all IDP ensembles.**

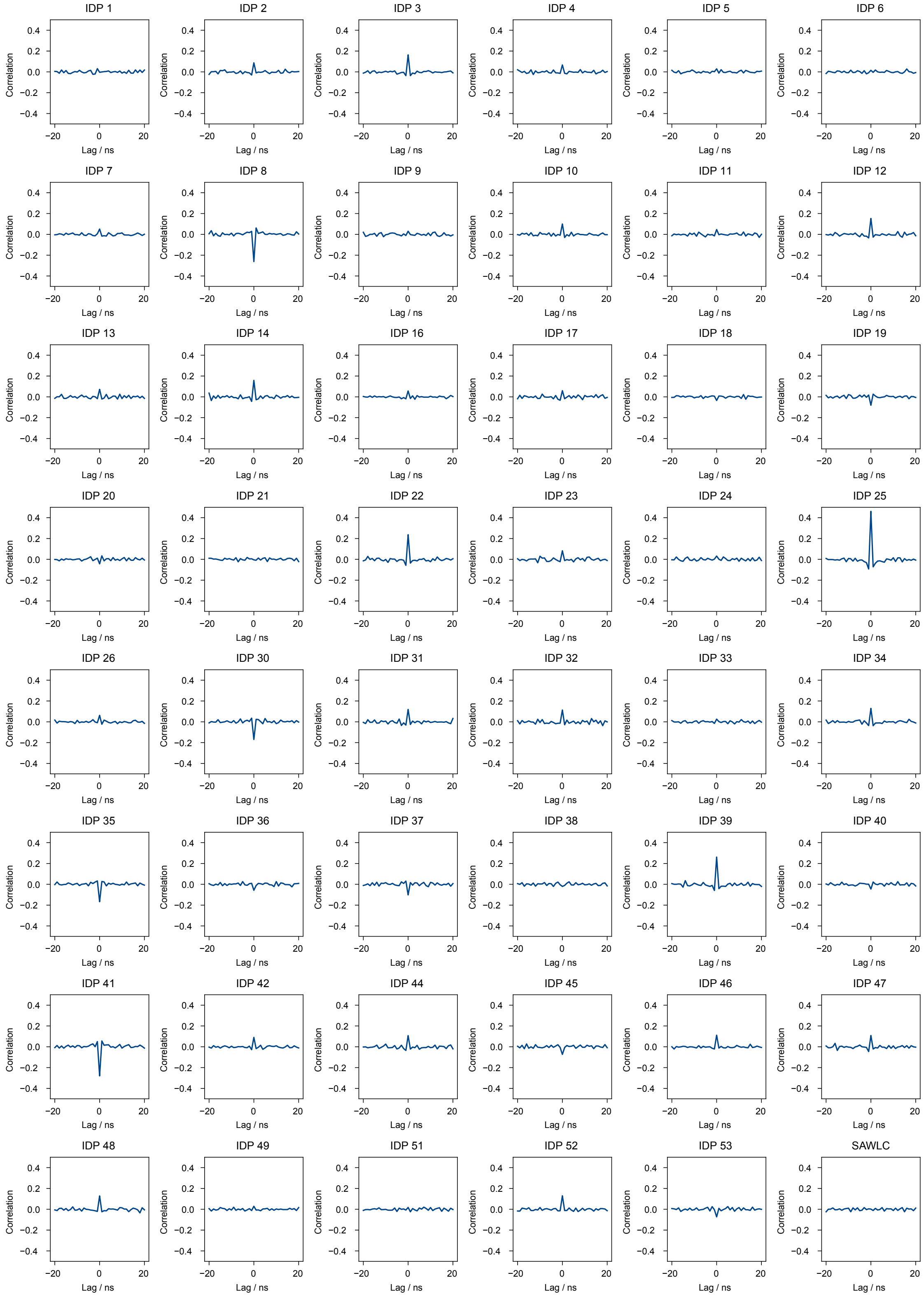


Figure S17: **Time-lagged cross-correlation between fluctuations in $\Delta R_\mathrm{g}(t)$ and local compactness asymmetry $\Delta\gamma_{R_\mathrm{g}}(t)$ for individual IDP ensembles.** Each panel represents one IDP.

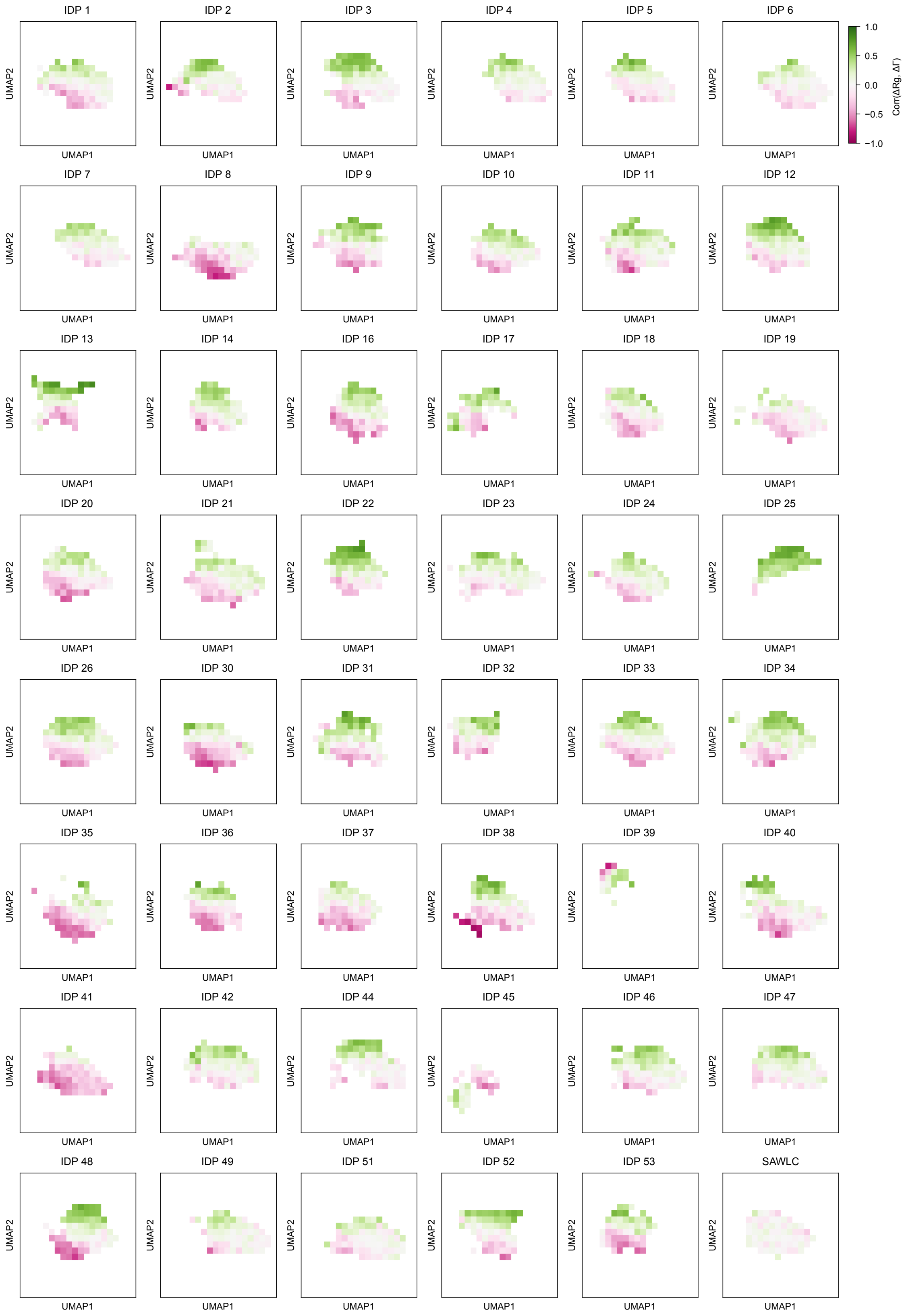


Figure S18: **Grid-wise cross-correlation at zero lag time ($\tau = 0$) between $\Delta R_g(t)$ and $\Delta\gamma_{R_g}(t)$ mapped onto the UMAP embeddings for individual IDP ensembles.** Each panel represents one IDP.

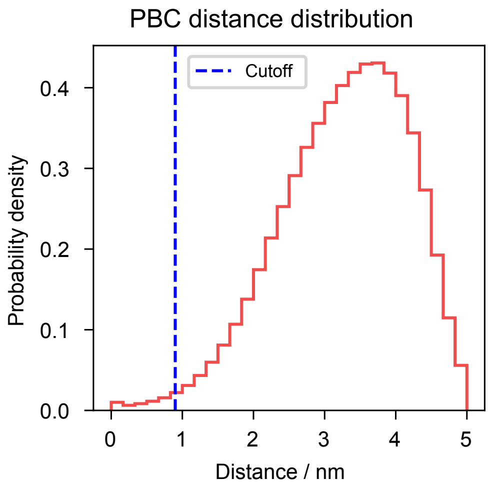


Figure S19: **Distribution of the minimum distances between atoms across periodic boundary condition (PBC) images.** The blue line indicates the 0.9 nm cutoff for nonbonded interactions. Notably, 99% of the structures have their minimum interatomic distance greater than this cutoff.

### References


(S1) Pearson, K. X. On the criterion that a given system of deviations from the probable in the case of a correlated system of variables is such that it can be reasonably supposed to have arisen from random sampling. *The London, Edinburgh, and Dublin Philosophical Magazine and Journal of Science* **1900**, *50*, 157–175.

(S2) Lalmansingh, J. M.; Keeley, A. T.; Ruff, K. M.; Pappu, R. V.; Holehouse, A. S. SOURSOP: A Python package for the analysis of simulations of intrinsically disordered proteins. *Journal of Chemical Theory and Computation* **2023**, *19*, 5609–5620.
